# Clonal Heterogeneity in Human Pancreatic Ductal Adenocarcinoma and Its Impact on Tumor Progression

**DOI:** 10.1101/2025.02.11.637729

**Authors:** Despoina Kalfakakou, Daniel C. Cameron, Emily A. Kawaler, Motoyuki Tsuda, Lidong Wang, Xiaohong Jing, Cristina Hajdu, Dylan L. Tamayo, Yoona Shim, Amanda Ackermann, Daniel Weissinger, Hayley Zimny, Rosmel Hernandez, Matthew Beier, Dacia Dimartino, Peter Meyn, Kalina Rice, Shanmugapriya Selvaraj, Cynthia Loomis, Adriana Heguy, Amanda W. Lund, Rosalie C. Sears, Theodore H. Welling, Igor Dolgalev, Aristotelis Tsirigos, Diane M. Simeone

## Abstract

Pancreatic ductal adenocarcinoma (PDAC) is an aggressive cancer characterized by profound desmoplasia and cellular heterogeneity, which cannot be fully resolved using traditional bulk sequencing approaches. To understand the contribution of this heterogeneity to PDAC biology, we analyzed a large cohort of primary human PDAC samples (n = 62), profiling 443,451 single cells and 53,236 spatial transcriptomic spots using a combined single-cell RNA sequencing and spatial transcriptomics approach. Our analysis revealed significant intratumoral heterogeneity, with multiple genetically distinct neoplastic clones co-existing within individual tumors. These clones exhibited diverse transcriptional states and subtype profiles, challenging the traditional binary classification of PDAC into basal and classical subtypes; instead, our findings support a transcriptional continuum influenced by clonal evolution and spatial organization. Additionally, these clones each interacted uniquely with surrounding cell types in the tumor microenvironment. Phylogenetic analysis uncovered a rare but consistent classical-to-basal clonal transition associated with *MYC* amplification and immune response depletion, which were validated experimentally, suggesting a mechanism driving the emergence of a more aggressive basal clonal phenotype. Spatial analyses further revealed dispersed clones enriched for epithelial-to-mesenchymal transition (EMT) activity and immune suppression, correlating with metastatic potential and colonization of lymph node niches. These dispersed clones tended to transition toward a basal phenotype, contributing to disease progression. Our findings highlight the critical role of clonal diversity, transcriptional plasticity, and TME interactions in shaping human PDAC biology. This work provides new insights into the molecular and spatial heterogeneity of PDAC and offers potential avenues for therapeutic intervention targeting clonal evolution and the mechanisms driving metastasis.

## INTRODUCTION

Pancreatic ductal adenocarcinoma (PDAC) is one of the most lethal human cancers, with an overall five-year survival rate of 13%^1^. This poor prognosis is attributed to several factors, including late diagnosis due to lack of early symptoms, a highly aggressive nature and early metastasis, and the ineffectiveness of standard chemotherapy and immunotherapy in most patients^2,3^. Notably, approximately 50% of new diagnoses present with metastatic disease; the average survival for these patients is less than one year^4^.

Genomic alterations and the immunosuppressive nature of the tumor microenvironment (TME) are critical drivers of PDAC progression and therapeutic resistance^5,6^. While key genomic drivers, including *KRAS*, *TP53*, *CDKN2A*, and *SMAD4*^7–11^, and discrete transcriptional subtypes (basal and classical)^12–15^ have been identified, the exact ways in which these features contribute to disease progression remain largely unknown. Among the transcriptional subtypes of PDAC, the basal subtype is associated with increased aggressiveness, characterized by enhanced epithelial-to-mesenchymal transition (EMT), immune evasion, and poor differentiation, all of which contribute to worse clinical outcomes. In contrast, the classical subtype demonstrates better differentiation, reduced EMT activity, and improved surviva_l_^16,17^. Despite the clinical relevance of these subtypes, the mechanisms driving their distinct phenotypes and the transitions between them remain poorly understood.

Although bulk sequencing and traditional methodologies have advanced our understanding of PDAC, they fail to capture the cellular and spatial heterogeneity that drives tumor progression and therapeutic resistance. Recent studies, including work by our group, have demonstrated significant intra- and intertumoral heterogeneity^18–25^ in human PDAC samples that bulk technologies failed to capture. These studies have underscored the extensive heterogeneity of PDAC tumors across transcriptional profiles, clonal populations, and TME composition, as well as the coexistence of cells exhibiting different subtypes within individual tumors, thereby highlighting the complex cellular architecture of this malignancy. However, the mechanisms by which clonal populations evolve, interact with their surroundings, and drive disease progression and metastasis remain poorly defined.

In this study, we address these key gaps in knowledge by applying single-cell and spatial transcriptomics to investigate the heterogeneity and clonal evolution of PDAC. We hypothesized that distinct clonal populations within tumors possess unique genomic and transcriptional features that shape their interactions with the TME, influencing disease progression and metastatic potential. To test this, we analyzed a cohort of 62 primary human PDAC tumor samples, freshly collected from 59 patients – the largest cohort studied to date – using scRNA-seq and spatial transcriptomics, uncovering novel features of intratumoral heterogeneity and mechanisms driving disease progression. Our analysis identified distinct clonal populations within individual tumors, characterized by unique copy number variations (CNVs), activation of specific signaling pathways, and subtype-specific transcriptional programs, as well as distinct interactions with the TME. We further explored clonal evolution, uncovering a specific event during which clones transition from the classical to the basal subtype, associated with amplification of the *MYC* locus and immune modulation. Spatial analysis revealed that dispersed clones exhibited elevated EMT activity, immune response depletion, and appeared to be transitioning toward a basal phenotype, correlated with metastatic potential to lymph nodes.

## RESULTS

### Multimodal profiling of human primary PDAC tumors

Our cohort comprises 62 primary PDAC tumor samples from 59 patients, including 6 pairs of matched samples collected before and after chemotherapy (Fig.1a). The PDAC specimens were obtained through surgical resection (n=24) or endoscopic ultrasound-guided core biopsy (n=38) of primary pancreatic lesions. 43 samples were collected prior to chemotherapy, while 19 were from patients who had received chemotherapy before sample collection (FOLFIRINOX-based n=12, gemcitabine/abraxane-based n=6, and mixed chemo-RT n=1).

A pancreatic pathologist assessed the histopathology of each tumor specimen included in the cohort to verify malignancy. Each tumor was then subjected to targeted gene panel sequencing to identify clinically relevant mutations, while 14 of the samples obtained via resection were additionally profiled with low-passage whole genome sequencing (WGS) to detect CNV events (Methods). After preparing single-cell suspensions, all samples included in the internal cohort were profiled with scRNA-seq using the 10x Genomics Chromium system (Fig. 1b, Methods). Furthermore, 11 samples were analyzed with 10x Genomics Visium FFPE spatial transcriptomics (ST). Among these, two tumor sections contained lymph nodes with metastasis, which were profiled in separate experiments, bringing the total number of ST experiments to 13.

**Figure 1.**
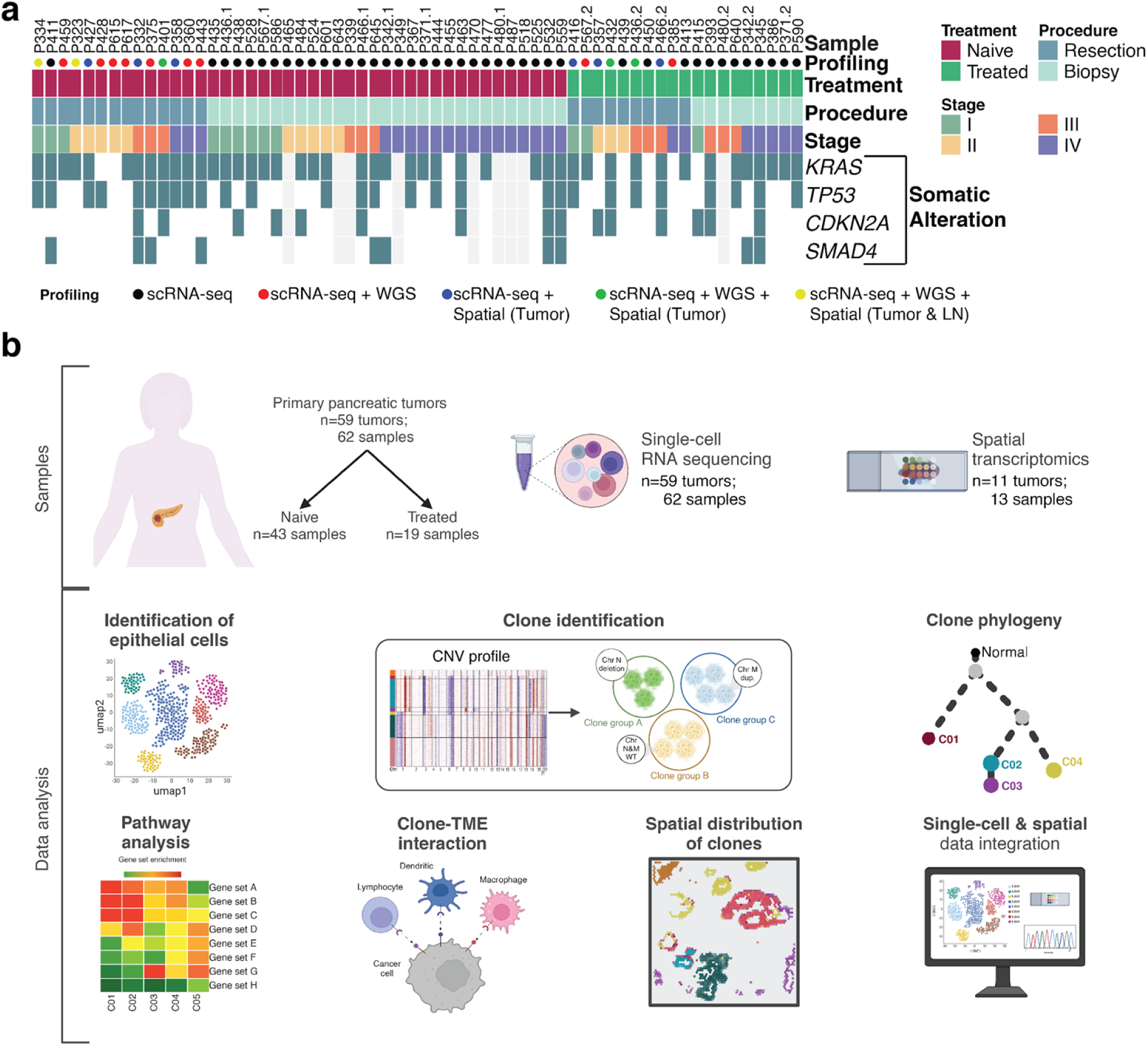
Mapping inter-tumoral heterogeneity across patient samples in PDAC with scRNA-seq. **a,** Cohort description and experimental design. **b,** Workflow depicting study design and analysis. Designed with BioRender©.

### The landscape of the PDAC TME at the single-cell level

Following standard pre-processing and quality filtering of the raw scRNA-seq data (Methods), 443,451 cells were available for further analysis. Using unsupervised graph-based clustering and visualization via Uniform Manifold Approximation and Projection (UMAP), we identified over 22 cell types (Ext. Fig. 1a,b), similar to what we had identified in a previous study^18^. The cell types included epithelial cells (including normal epithelial and malignant tumor cells), T cell subsets, NK cells, fibroblasts, endothelial cells, myeloid cell subsets, B/plasma cells, mast cells, and a subset of acinar cells. Focusing on the epithelial cell population, we observed that while cells from other populations clustered together irrespective of their sample origin, epithelial cells formed patient-specific clusters (Ext. Fig. 1c), indicating significant intertumoral or patient heterogeneity, a phenomenon that has been previously documented in both pancreatic cancer and other solid tumors^18,26,27^. The identified clusters were annotated using canonical markers and gene expression profiles (Ext. Fig. 1d-i, Methods). Differential analysis of the cell type composition between naïve and treated samples revealed no significant differences across most cell types, apart from a slight increase in the endothelial cell population in treated samples (false discovery rate (FDR) = 0.03) (Ext. Fig. 1k). While our cohort of 62 human PDAC samples provides substantial insight, it is possible that subtle compositional differences remain undetected due to the complexity of the dataset and the inherent variability of human PDAC samples.

### Subpopulations within the tumor epithelial compartment reveal a dynamic process of clonal evolution

While intertumoral heterogeneity in PDAC has been documented in several studies, intratumor heterogeneity remains less explored. To address this gap, we conducted a detailed analysis of the neoplastic epithelial cell compartment in each sample. Initial clustering of epithelial cells revealed the presence of distinct epithelial subpopulations within most tumor samples, indicating the potential coexistence of multiple genetic clones. Building on these observations, we hypothesized that the identified transcriptional subpopulations represent distinct clonal populations shaped by both shared and unique CNV events. To test this hypothesis, we applied inferCNV^28^ to analyze each sample individually, identifying CNVs across epithelial cells. To ensure robustness, we focused on samples containing more than 100 epithelial cells, resulting in a final study cohort of 50 samples. The majority of epithelial cells across all samples exhibited multiple CNV events, indicating malignancy and clearly differentiating malignant cells from CN-normal epithelial cells. (Figure 2a,f, Suppl. Fig. 1). Hierarchical clustering of the CNV profiles of individual cells enabled the identification of malignant cell clusters with shared CNV profiles, indicating distinct genetic clones. Clusters were classified as separate clones based on distinct CNV profiles, identified through manual inspection of chromosomal alterations. We then constructed a phylogenetic tree for each patient sample using the inferred CNVs of each clone, based on the principle that descendant clones accrue additional CNVs absent in their predecessors. When two or more clones exhibited distinct sets of additional CNVs, we inferred the presence of an unobserved common ancestor from which these clones diverged prior to acquiring their unique CNVs (Fig. 2b,g, Suppl. Fig. 1).

**Figure 2.**
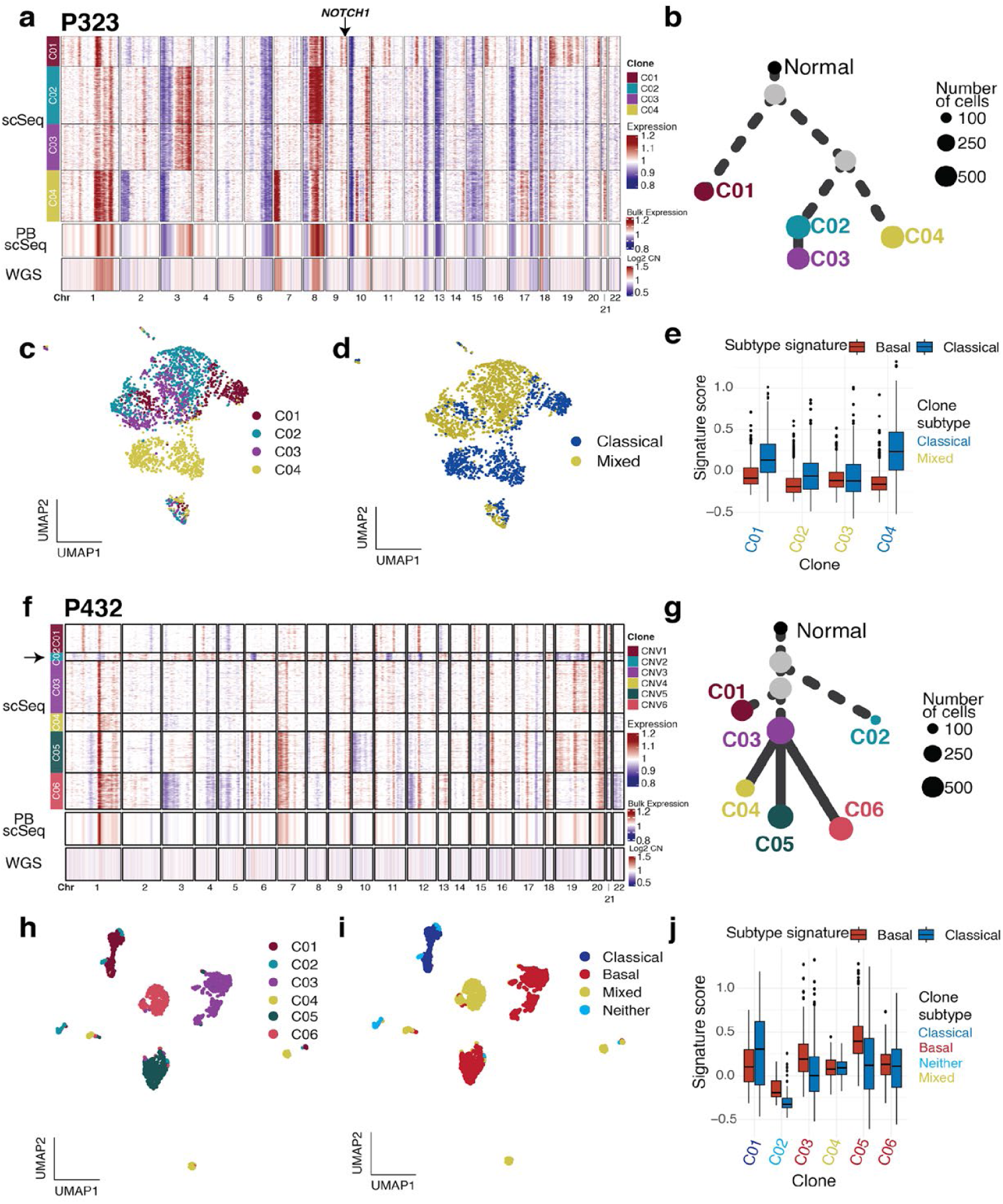
Clonal composition and subtype heterogeneity of malignant epithelial cells. **a,** CNVs inferred from single-cell RNA sequencing (scSeq) data (top), pseudo bulking (BP) of the same scSeq data (middle), and matched whole genome sequencing (WGS - bottom) of a representative sample (P323). **b,** Inferred phylogenetic tree for P323. **c,** UMAP of malignant epithelial cells of P323 color-coded by clone. **d,** UMAP of malignant epithelial cells of P323 color-coded by subtype per clone. **e,** Boxplots of subtype scores per clone for P323. **f,** CNVs inferred from single-cell RNA sequencing (scSeq) data (top), pseudo bulking (BP) of the same scSeq data (middle), and matched whole genome sequencing (WGS - bottom) of a representative sample. **g,** Inferred phylogenetic tree for P432. **h.** UMAP of malignant epithelial cells of P432 color-coded by clone. **i,** UMAP of malignant epithelial cells of P432 color-coded by subtype per clone. **j,** Boxplots of subtype scores per clone for P432.

Identifying cell clusters with shared CNV profiles revealed genetic differences between distinct clones within the same tumor. For example, in patient sample P323, a representative sample collected before chemotherapy treatment, we identified a unique clone (C01) with a chromosome 9q34 amplification, containing *NOTCH1* (Fig. 2a), a gene linked to cell proliferation and increased metastatic potential. Notably, WGS of this sample did not detect all the CNVs identified through scRNA-seq, including the 9q34 amplification, in contrast to other CNVs that were confirmed; this limitation is likely due to the presence of the amplification in only a single clone. Indeed, pseudo-bulking of the normalized expression output of inferCNV shows how the signal of some CNVs can be attenuated or completely lost, underscoring the sensitivity of scRNA-seq in detecting CNVs (Fig.2a). Projection of clones onto the UMAP of malignant epithelial cells, calculated based on the gene expression profiles of individual cells, revealed two patterns: in some samples, cells within a single clone formed distinct clusters, while in others, cells from multiple clones clustered together. For instance, in sample P323, clones C02 and C03 -which differ only by a single deletion on chromosome 15-appeared to cluster together (Fig. 2c), suggesting that while genetic divergence has begun, it has not yet significantly impacted their transcriptional profiles (Fig. 2b). These scenarios suggest that clones with slightly differing CNV profiles will exhibit similar gene expression patterns.

Given the association of the basal subtype with increased aggressiveness and poor clinical outcomes in PDAC, we next sought to investigate how intratumoral subtype heterogeneity might contribute to PDAC progression. Classifying each clone based on the dominant enriched transcriptional subtypes^12^ revealed that in P323, two out of four clones predominantly consisted of classical cells (Fig. 2d, e). For clones C02 and C03, the population consisted of a mix of classical cells and cells that did not predominantly align with either the basal or classical signature, which we refer to as “mixed”.

In patient sample P432, which was collected post-chemotherapy through surgical resection, we observed distinct genomic events across the clones, with one small clone (C02) exhibiting significantly more diversity compared to the others (Fig.2f). Notably, P432 was the only sample where the CNVs identified through scRNA-seq were not fully confirmed by WGS, highlighting the limitations of WGS in detecting certain small or subclonal events in genetically diverse tumors (Fig.2f, Suppl. Fig. 1). The construction of a phylogenetic tree for this sample reveals that C02 diverged early in the tumor’s evolutionary history, while C04, C05, and C06 share C03 as a common ancestor (Fig. 2g). The clones in this sample form distinct transcriptional clusters, suggesting enrichment in different signaling pathways that may be driven by their genetic profiles (Fig.2h). In contrast to patient sample P323 that was predominantly classical, here the classification of clones by their dominant subtype reveals the coexistence of both classical (C01) and basal (C03, C06) clones within a single tumor, as well as the presence of both basal and classical cells within individual clones (C04, C06) (Fig.2i,j). Additionally, the small, genetically diverse clone (C02) did not predominantly align with either the basal or classical subtypes and was thus classified as “neither”.

Our findings demonstrate significant intratumoral heterogeneity in PDAC, with distinct genetic clones co-existing within individual tumors, highlighting the complex and dynamic nature of tumor evolution. Furthermore, we observed the concurrent presence of classical and basal subtypes in many tumors, challenging the traditional binary classification of PDAC. Instead, our findings emphasize that diverse clonal populations contribute to a diversity of transcriptomic signatures within tumors, underscoring the complexity of PDAC biology within each individual patient’s tumor.

### Characterizing the clonal intra- and intertumoral heterogeneity in PDAC

Building on our identification of distinct clones within tumors, we sought to characterize the transcriptional and genetic diversity of these clonal populations. We hypothesized that the genomic differences defined by CNVs influence the transcriptional programs of clones and contribute to the basal and classical subtype classifications. To test this, we examined the relationship between the genomic events defining the clones and their predominant subtypes, as well as how the genomic profiles of these clones correlate with the activation of key cancer-related pathways.

Our analysis revealed an average of 4.8 clones per patient (range: 2–9; standard deviation (SD): 1.9; standard error of the mean (SEM): 0.26), while the number of cells in each clone averaged 568 cells (range: 21–7372; SD: 900; SEM: 58) (Fig. 3a). Although samples obtained via core biopsy had fewer epithelial cells than the samples from resections (Suppl. Fig.2a), we did not find any significant difference in the number of clones between the two groups (Suppl. Fig.2b).

**Figure 3.**
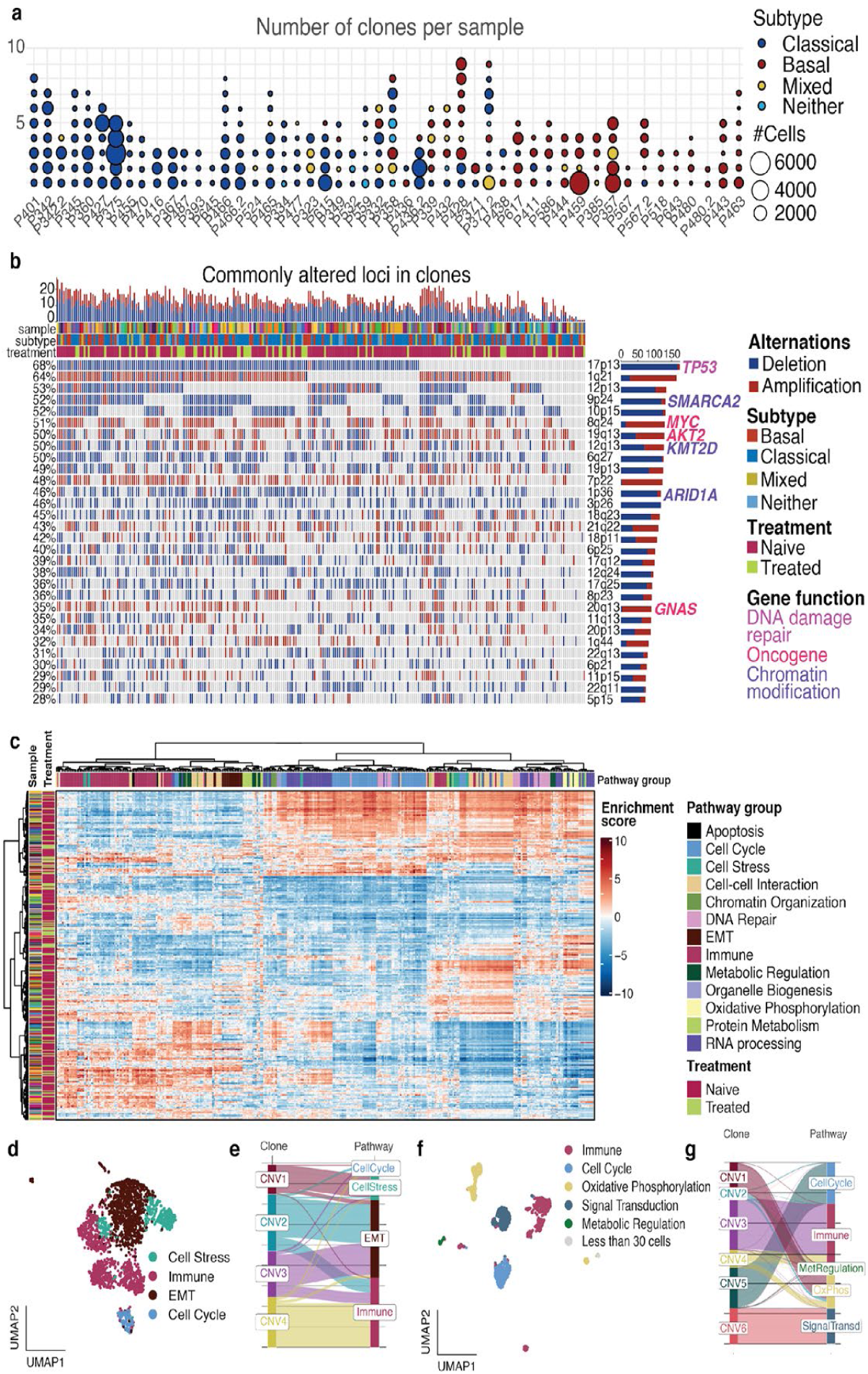
Basic characterization of the clonal heterogeneity landscape. **a,** Dot plot of number of clones per sample showing the number of cells and the major subtype of each clone. **b,** Loci altered in more than 25% of clones. **c,** Differential pathway enrichment of tumor transcriptional clusters defined after unsupervised clustering of the malignant epithelial cells of each patient sample. Columns are clusters and rows are different pathways. d, UMAP of malignant epithelial cells of P323 color-coded by pathway group per transcriptional cluster. **e,** Alluvial plot showing the correlation between the pathway expressed by each cluster and the cells in each clone for sample P323. **f,** UMAP of malignant epithelial cells of P432 color-coded by pathway group per transcriptional cluster. **g,** Alluvial plot showing the correlation between the pathway expressed by each cluster and the cells in each clone for sample P432.

To classify all of the clones in each tumor sample into subtypes, we used the Moffitt subtype signature expression. Previous analysis on the bulk-derived classifications by Collisson, Bailey, and Puleo^13–15^ revealed that the Moffitt-basal signature overlaps with the Collisson-quasi-mesenchymal, Puleo-pure-basal, and Bailey-squamous signatures, while the Moffitt-classical signature aligns with the Collisson-classical, Puleo-pure-classical, and Bailey-progenitor signatures, with no strong expression observed for other signatures described in these studies^18^. Therefore, for the remainder of this manuscript, we use the Moffitt-classical and Moffitt-basal subtypes to describe the two major subtypes (Suppl. Fig. 2c,d). A small subset of cells (approximately 5%) tested present in most samples (41/50), had negative scores for both basal and classical signatures, lacking enrichment of either of the signatures, and were thus classified as ‘neither’ (Suppl. Fig. 2d).

The identified clones, however, were predominantly of the classical (n=134) or basal subtype (n=75). Sixteen of the samples (32%) were predominantly classical, eight samples (16%) were predominantly basal, and twenty-six (52%) contained a mixture of at least two clones from each subtype (Fig. 3a, Suppl. Fig. 2e).

We then analyzed the CNVs identified in each clone. In our cohort, the most frequently altered locus was 17p13, containing the TP53 gene, which was part of a CNV in 174 out of 241 clones (68%); among these alterations, 98% involved a deletion, as expected^10,29^. Another commonly altered locus was 8q24, which includes the MYC gene, with gains observed in 115 clones (44.9%) and deletions in 15 clones (5.9%). Additional common alterations were observed in loci harboring other genes previously implicated in pancreatic cancer^10,29^, including 9p24 (*SMARCA2*; 52% of clones), 19q13 (*AKT2*; 50% of clones), 12q13 (*KMT2D*; 50% of clones), 1p36 (*ARID1A*; 46% of clones), and 20q13 (*GNAS*; 35% of clones) (Fig. 3b).

When examining the CNV differences between basal and classical clones, we observed that the loci altered included genes defining the basal and classical subtype signatures^12^. Specifically, 21q22, which includes classical signature genes *TFF1* & *TFF2*, was significantly amplified in classical clones and deleted in basal clones. Furthermore, loci 1q21 and 17q21, which include the basal subtype signature genes *S100A2* and *KRT15* & *KRT17*, respectively, were significantly amplified in basal clones (Suppl. Fig. 2f).

In order to investigate whether there is a link between the genetic and transcriptional profiles of the clones, we performed pathway analysis. First, we identified transcriptional clusters in the malignant epithelial cells of each sample using unsupervised, graph-based clustering. These clusters were then used for pathway enrichment analysis with gene set enrichment analysis (GSEA). Hierarchical clustering of these pathways revealed correlations between specific pathway groups (Fig. 3c); for example, pathways related to cell proliferation and oxidative phosphorylation were frequently co-enriched within the same transcriptional clusters, as were immune response and EMT-related pathways (Suppl. Fig. 2g). While one might expect treatment status to affect enriched pathways, we saw no evidence of that here (Fig. 3c). Projection of the top pathway groups that were enriched in each transcriptional cluster onto the UMAP of malignant epithelial cells from each sample revealed a notable association between the pathways enriched in each transcriptional cluster and the corresponding clone cells. This observation suggests that the genetic profile of a clone may influence the transcriptional phenotype of the cells. For instance, in sample P323 (Fig. 2c, 3d,e), clone C01 is predominantly composed of cells enriched in stress-related pathways, while clone C04 mainly consists of cells enriched in immune-related pathways. Pathways associated with cell cycle activity are enriched across cells from multiple clones in this sample, suggesting that cell cycle regulation, essential for tumor growth, may be driven by shared oncogenic pressures rather than strictly by CNV profiles. In contrast, in sample P432 (Fig. 2h, 3f,g), cell cycle activity is restricted to a subset of cells within clone C05, suggesting that in some cases, microenvironmental cues or specific genomic or epigenomic alterations may play a more dominant role. These observations highlight the dual influence of tumor-wide oncogenic pressures and clone-specific factors in regulating cell cycle activity within PDAC tumors. Overall, our results suggest that subtype heterogeneity in human PDAC tumors is driven by distinct genetic events, as demonstrated by the amplification or deletion of loci encoding key marker genes associated with these subtypes. Pathway enrichment analysis further reveals that the CNV profiles of clones shape their transcriptional phenotypes, driving both subtype differentiation and the activation of key pathways.

### Clonal diversity shapes a heterogeneous landscape of interactions with the PDAC TME

Building on our observations of clonal heterogeneity in genetic events, subtypes, and transcriptional programs, we hypothesized that individual clones may also differ in how they interact with other cells within the TME. To identify how malignant cell clonal evolution impacts the TME in PDAC, we performed a ligand-receptor interaction (LRI) analysis between each clone and different TME cell types. Our analysis revealed that different clones interact with TME cell types in distinct ways, even within the same sample (Ext. Fig. 2a). Importantly, this result was not influenced by the size of the clone, as several clones with a large number of cells exhibited a low number of interactions with other cell types and vice versa (Ext. Fig. 2a).

To quantify interaction heterogeneity, we categorized clones into three groups based on the number of interactions they exhibited with each cell type: low, intermediate, or high. Next, each patient sample was classified based on the overall interaction profile of its clones with other TME cell types. Samples were categorized as having low TME interactions if all clones were in the low or intermediate interaction groups, intermediate TME interactions if all clones were in the intermediate group, and high TME interactions if all clones were in the high or intermediate groups. In cases where at least one clone exhibited low interactions and another clone displayed high interactions with the same cell type, the sample was classified as displaying heterogeneous interactions with this cell type (Ext. Fig. 2b). Of the 50 tumors analyzed, 37 (74%) displayed heterogeneous interactions with at least one cell type in theTME, a pattern that appeared independent of treatment or even the number of clones within a sample (Ext. Fig. 2b). Among the remaining samples, three displayed only low numbers of interactions with other cells in the TME, while one showed consistently high interactions across all its clones with all the cell types. The rest of the samples exhibited varying numbers of interactions between their clones and different cell types.

Heatmaps for representative six samples (Suppl. Fig. 3a-f) illustrate the interaction counts between clones and key TME cell types, along with clone sizes and cell type frequencies. These heatmaps highlight how samples can uniformly fall into the low (Suppl. Fig. 3a), intermediate (Suppl. Fig. 3b) or high (Suppl. Fig. 3c) interaction groups or exhibit distinct heterogeneity patterns (Suppl. Fig. 3d-f). For instance, some clones show strong interactions with CD8+ T cells or fibroblasts, while others within the same sample interact minimally with these cell types, revealing variability in interaction profiles.

Building on our observations of clonal heterogeneity in LRIs within theTME, we sought to determine whether CNVs contribute to this variability by influencing ligand and receptor availability. To systematically assess the impact of CNVs on cell-cell interactions, we calculated a CNV-weighted interaction score for each ligand-receptor pair (Methods). This score accounts for how frequently a gene is amplified or deleted across clones and how often its interactions are significantly detected. Specifically, for amplifications, the score reflects the number of times an amplified gene participates in a significant interaction, while for deletions, it quantifies the number of times the interaction is absent when the gene is deleted (Ext. Fig. 2c,d).

Our analysis reveals that CNV alterations play a pivotal role in shaping LRIs within the TME, influencing immune evasion, stromal remodeling, and potential therapeutic strategies. Amplifications of the *PVR* and *NECTIN2* locus (19q13), which is one of the most commonly amplified loci (Fig. 3b) drive enhanced immune checkpoint interactions between the two molecules and *TIGIT* (Ext. Fig. 2c), suppressing T cell and NK cell cytotoxicity^30,31^. This suggests that TIGIT inhibitors would offer a more promising therapeutic avenue than PD-1/PD-L1 inhibitors, especially given the observed deletions of *PDCD1LG2* and *CD274*, which suppress PD-1/PD-L1 interactions (Ext. Fig. 2d) and diminish their activity as immune checkpoints^18,32^.

In tumor-associated macrophages (TAMs), *AXL* amplification enhances GAS6-AXL signaling (Ext. Fig. 2d), driving immune suppression and stromal remodeling, and is linked to therapy resistance and poor prognosis^33^. Stromal interactions are driven by amplified *PDGFA* promoting PDGFA-PDGFRA signaling (Ext. Fig. 2c). PDGFA-PDGFRA activation has been shown to reprogram the tumor stroma, contributing to an immunosuppressive TME and enhancing tumor invasiveness through fibroblast activation and extracellular matrix (ECM) remodeling, while dual blockade of PDGFRα/β has been proposed as a strategy to reverse these stromal effects and improve the efficacy of immune checkpoint inhibitors in fibrotic cancers ^34^.

Our findings demonstrate that LRIs are not uniform across clones, with some interactions being enhanced or suppressed due to CNV-driven alterations. This raises the question of whether these patterns are consistent across all clones or if additional factors contribute to interaction heterogeneity beyond CNV effects. To explore this further, we focused on interactions between clones and CD8+ T cells mediated by checkpoint molecules, since checkpoint inhibitors have yet to demonstrate efficacy in PDAC clinical trials, apart from a small subset (1%) of mismatch repair deficient tumors^35,36^. While some clones exhibited strong interactions with CD8+ T cells, others within the same sample showed no interactions at all (Ext. Fig. 2e). Notably, most clones demonstrated significant interaction scores through the PVR-TIGIT axis, whereas no clones in any sample interacted with CD8+ T cells via the PDCD1LG2-PDCD1 axis (Ext. Fig. 2e). These patterns of interaction heterogeneity highlight the complexity of immune evasion in PDAC and underscore the challenges of effective immunotherapy.

This analysis reveals significant heterogeneity in the interactions between malignant clones and different cell types within the TME, independent of clone size. This variability is particularly evident in interactions with immune cells, such as CD8+ T cells, posing challenges for immunotherapy in PDAC. However, our findings also point to potential therapeutic opportunities by targeting CNV-driven interactions, such as the PVR or NECTIN2 interaction with TIGIT or the GAS6-AXL axis, where *AXL* is frequently amplified. By leveraging the differential contributions of amplified and deleted genes within the TME, these insights provide a framework for prioritizing targeted interventions that may help overcome the limitations of current immunotherapies to effectively treat PDAC.

### Patterns of clonal expansion and elimination after treatment with chemotherapy

To assess the impact of chemotherapy on clonal evolution in PDAC, we analyzed matched pre- and post-treatment samples from six patients. Across all cases, we observed distinct patterns of clonal expansion and elimination.

As an example, in sample P466, three clones were retained post-treatment (C02, C06, C11), five clones were eradicated (C01, C05, C07, C08, C09), and three new clones emerged (C03, C04, C10) (Ext. Fig. 3a-e). Notably, C07, C08, and C09 formed an entire branch of the phylogenetic tree and harbored a unique 12q13 amplification absent in other clones. However, this amplification was not consistently associated with clone eradication in other samples. Although clones C02 and C06 in this sample constituted a small percentage of tumor cells before treatment, they expanded markedly post-treatment. In contrast, C11, which also persisted after therapy, maintained a small population comparable to its pre-treatment levels (Ext. Fig. 3c,e).

LRI analysis revealed substantial heterogeneity in how clones interacted with the TME before and after treatment. For example, while both C02 and C06 of patient sample P466 expanded after treatment, they displayed distinct interaction patterns (Ext. Fig. 3f,g). Before treatment, C02 had limited interactions with other cells but displayed increased interactions with all TME cell types after therapy. In contrast, C06 initially exhibited strong interactions with T cells and myofibroblastic cancer-associated fibroblasts (myCAFs), which diminished following treatment. However, its interactions with the myeloid compartment remained stable, while those with inflammatory CAFs (iCAFs) and endothelial cells increased post-treatment (Ext. Fig. 3f,g).

While patterns of engagement with most TME components varied across clones, increased interactions with iCAFs and endothelial cells were consistently observed in clones retained post-treatment, whether they expanded or remained stable in size (Ext. Fig. 3g-k). iCAFs are known to promote a pro-inflammatory microenvironment while simultaneously suppressing anti-tumor immune responses^37^, creating a supportive niche for resistant clones. Similarly, increased interactions with endothelial cells, key players in angiogenesis and vascular remodeling^38^, suggest an adaptive mechanism favoring clonal survival and potential progression under treatment-induced stress.

Together, these findings demonstrate that chemotherapy not only reshapes the clonal landscape of PDAC but also influences how surviving clones engage with the TME. The consistent increase in interactions with iCAFs and endothelial cells suggests that resistant clones adapt to treatment-induced stress by leveraging supportive stromal niches, highlighting potential targets for therapeutic intervention.

### MYC amplification enables a unique clonal transition from classical-to-basal phenotype

To gain a greater understanding of the landscape of clonal evolution in the entire patient cohort, we constructed a phylogenetic “forest”, consisting of sample-specific phylogenetic trees, based on their CNV profiles (Fig. 4). Color-coding the clones in each phylogenetic tree according to the predominant subtype of cells within each clone allowed tracking of clonal evolution patterns in each tumor, revealing significant intratumoral heterogeneity. While a large subset of trees (n=23) consisted primarily of classical clones (denoted with blue), either entirely or predominantly (all but one clone being classical), the presence of basal clones in these trees suggests the potential for evolutionary shifts toward a more aggressive phenotype. A smaller subset of trees (n=11) consisted primarily of basal clones (denoted with red), while the remaining trees contained both classical and basal clones, or clones without a predominant subtype (mixed, yellow) or that weren’t enriched in either the classical or basal signature (neither, light blue) (Fig. 4), once again underscoring the substantial heterogeneity present in PDAC and its potential role in driving tumor progression.

**Figure 4.**
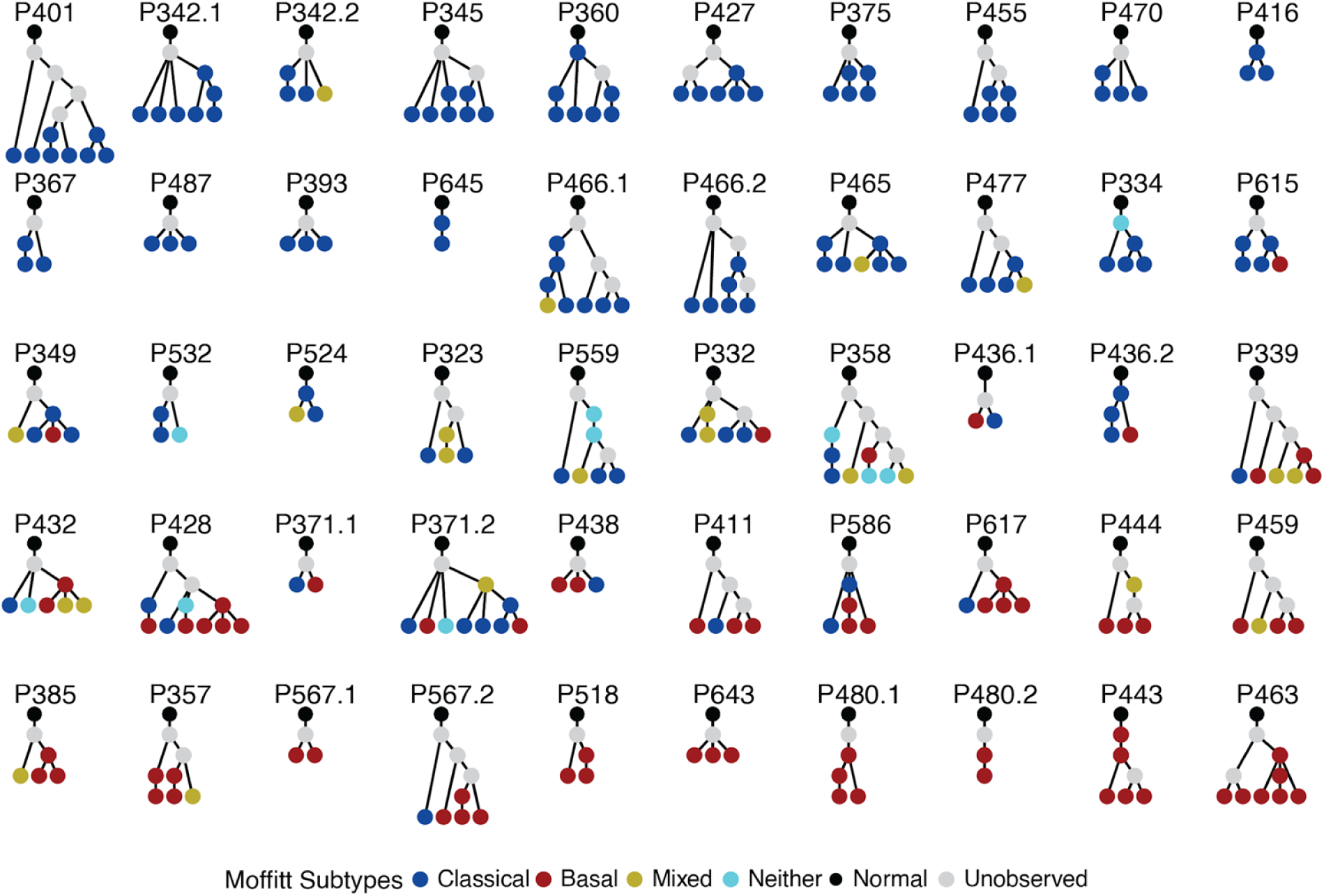
Phylogenetic forest reveals distinct patterns of evolution. Phylogenetic tree forest of patient tumor samples, constructed based on the CNV profiles of individual clones

Examination of the evolutionary paths of each tree revealed that both classical and basal clones typically preserve their subtype throughout evolution. Intriguingly, we detected an evolutionary transition from a classical to a basal clone in six samples; in contrast, we observed no instances of a basal to classical transition (Fig. 5a). In rare instances, we observed clones exhibiting neither subtype giving rise to classical and/or basal clones, such as the founder clone in sample P334 (second row, second from right, Fig 4). Conversely, classical and basal clones occasionally gave rise to progeny with mixed or neither subtype, highlighting the inherent plasticity of PDAC tumors despite most clones being predominantly classical or basal.

**Figure 5.**
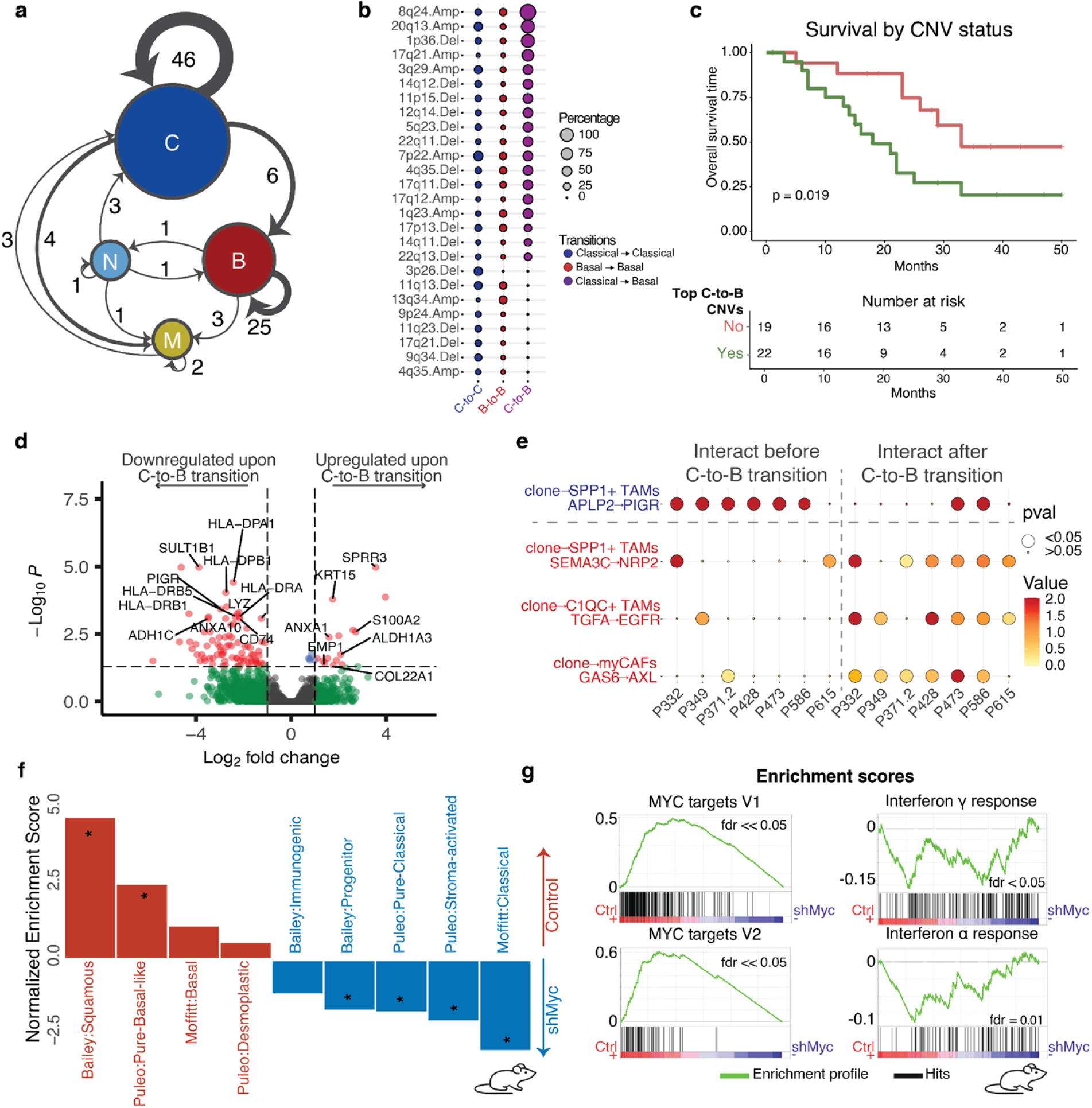
Subtype transitions in PDAC evolution. **a,** Network diagram illustrating subtype evolutionary transitions. **b,** Percentage of commonly altered loci for the most common evolutionary transitions. **c,** Survival analysis of patients with or without one of the classical-to-basal transition CNVs. **d,** Genes differentially expressed upon the classical to basal transition. Plotted are all enriched signatures, with the ones with an FDR<0.05 being denoted by an asterisk. **e,** Ligand-receptor interactions between cancer cells and various compartments in the TME, comparing clones in the classical-to-basal transition pairs. **f,** *Myc* knockdown in KMC-derived cell lines promotes a basal to classical transition (red : KMC control; blue: KMC shMyc). **g,** *Myc* knockdown in KMC cell lines results in enrichment of immune-related pathways.

To identify the genetic basis underlying the examples of observed classical-to-basal transition (Fig. 5a), we examined the CNVs that differed between the ancestral and their descendant clones. In all cases, we identified an acquisition of an amplification in 8q24, a locus containing the MYC gene, suggesting that it may be a critical driver of the process. *MYC* amplification is common in many cancers and has been shown to contribute to tumor growth and metastasis in PDAC^11,39^. While amplification of 8q24 was seen in other clones as well, it occurred at a significantly higher frequency in the cases of classical-to-basal transitions (Fig. 5b). Other CNVs frequently associated with this transition included amplifications at the 17q21 and 20q13 loci and a deletion at 1p36. Notably, 17q21 harbors keratin genes, such as *KRT15*, *KRT16*, and *KRT17*, which are related to the basal signature^12,15^. These observations indicate that the classical-to-basal transition may be associated with a combination of key CNV events, with 8q24 amplification playing an important role.

To assess the clinical impact of these CNVs, we performed a survival analysis, comparing patient tumors harboring the aforementioned CNVs to those without. The analysis revealed that patients with tumors containing a CNV from the classical-to-basal transition signature had significantly worse overall survival than those without these CNVs (p = 0.019) (Fig. 5c).

Since 8q24 amplification was consistently gained during transitions, including classical-to-classical transitions even at a lower rate, we hypothesized that while it serves as a necessary driver of the classical-to-basal transition, additional molecular changes may be required to fully facilitate this shift. To explore this possibility, we performed differential gene expression and LRI analysis on the clone pairs participating in the classical-to-basal clonal transitions. Differential gene expression analysis (DGEA) showed that the clones that transition from a classical to a basal phenotype are marked by upregulation of genes that promote EMT, such as *S100A2*, *COL22A1*, *EMP1*, and *EGFR*, and downregulation of immune response-related genes, such as *PIGR*, *CD74*, *CXCL2*, and various HLA genes, particularly class II genes (Fig. 5d). On the contrary, clones that preserve the classical phenotype in the presence of 8q24 amplification, did not exhibit downregulation of the same signaling pathways during the transition (Suppl. Fig. 4a,b). These findings suggest that immune evasion may play a complementary role with genetic events in driving the classical-to-basal subtype transition.

To determine whether clones that underwent a classical-to-basal transition exhibit differential interactions with the TME, we performed LRI analysis on the clone pairs involved in these transitions. This analysis revealed a specific interaction between classical clones (prior to the transition) and SPP1+ TAMs, as well as three distinct interactions between basal clones (following the transition) and other cells within the TME (Fig. 5e). The interaction between classical clones and *SPP1*+ TAMs was facilitated via the APLP2-PIGR pathway, with PIGR standing out due to its decreased expression during the classical-to-basal transition. PIGR’s involvement in this pathway suggests a potential role in facilitating immune surveillance, as it has been implicated in tumor-immune interactions through processes like the transcytosis of dimeric IgA^40^. Basal clones emerging from these transitions established interactions in pathways associated with tumor progression and immune evasion, including the SEMA3C-NRP2 pathway with *SPP1*+ TAMs, the TGFA-EGFR pathway with *C1QC*+ TAMs, and the GAS6-AXL axis with myCAFs^41–44^. These findings suggest that during classical-to-basal clonal transition, malignant cells enhance interactions with tumor-promoting TAM subsets and stromal cells through pathways known to drive tumor progression and immune suppression, while concurrently downregulating immune-related genes, potentially facilitating immune evasion and the emergence of a permissive tumor microenvironment.

To test our hypothesis that MYC amplification is necessary to drive evolutionary transition from the classical to the basal subtype, we knocked down MYC expression in two independently derived tumor cell lines from a PDAC mouse model with Ptf1a-CreERT2-induced expression of constitutive deregulated MYC and mutant KRAS (KMCE)^45^, using doxycycline-inducible lentiviral shMyc (Methods).

To confirm whether *Myc* knockdown induces a reversal from a basal to a classical phenotype, we conducted GSEA using all published subtype signatures for PDAC, rather than only the Moffitt signature, as there is no direct correspondence between these gene signatures in mouse and human. In the KMCE PDAC cell lines basal subtype signatures were predominantly enriched, with the Bailey squamous and Puleo Pure-Basal signatures showing the strongest enrichment (FDR < 0.05) (Fig. 5f, red bars). In contrast, following *Myc* knockdown, the cells shifted to a classical subtype, with the Moffitt Classical, Puleo Pure-Classical, and Bailey progenitor signatures being most prominent (Fig. 5f, blue bars, Suppl. Fig. 4c,d). These results support that *Myc* knockdown drives a shift from a basal to a classical phenotype, and reinforce the role of *MYC* in driving the transition from the classical to a more aggressive basal phenotype in human PDAC clones.

To investigate whether the immune escape observed in human PDAC clones that underwent classical-to-basal transition is recapitulated in murine PDAC cells, we performed pathway enrichment analysis in KMCE cell lines with and without *Myc* knockdown. Consistent with the findings from human PDAC clones, *Myc* knockdown resulted in the enrichment of immune-response related pathways (Fig. 5g). This finding suggests that MYC amplification is a prerequisite for the transition from a classical to a basal phenotype. Additionally, the restoration of immune pathway activity upon MYC loss suggests that MYC plays an immunosuppressive role.

### Spatial distribution of tumor clones in pancreatic cancer

Having characterized clonal heterogeneity at the single-cell level, we next sought to understand how these clones are organized within the tumor microenvironment. To investigate the spatial distribution of clonal heterogeneity in PDAC tumor tissue, we profiled a subset (11 samples) from our scRNA-seq cohort using 10x Genomics Visium. Following quality control and filtering, we retained 53,236 high-quality spots from primary pancreatic tumors for analysis. Cell type deconvolution was conducted to identify the cellular composition within each spot, using the scRNA-seq data as a reference (Ext. Fig. 4a,b). Manual annotation of cell types, informed by unbiased clustering and marker gene expression, demonstrated a strong correlation with the deconvolution-based annotation, supporting the accuracy of our methodology (Ext. Fig. 4c,d). CAFs were enriched throughout the tissue (Ext. Fig. 4a,b,e); notably, while myCAFs were abundant across all samples, iCAFs were specifically enriched in the treated samples (Ext. Fig. 4e).

Given the observed mix of epithelial and other cell types, predominantly stromal cells, with no distinct regions of epithelial cells, we identified clones in our spatial data by focusing on spots with high estimated epithelial content, based on the deconvolution results. For each sample, a pathologist annotated regions containing normal epithelial cells and spots with malignant tumor cells (Fig. 6a). Using inferCNV, we analyzed spots with more than 35% epithelial content using the spots with less than 10% epithelial content for reference. Notably, we identified the same CNVs in both the spatial transcriptomics and their matched scRNA-seq samples, and there was overall concordance at the clone level (Fig. 6b,c, Suppl. Fig. 5).

**Figure 6.**
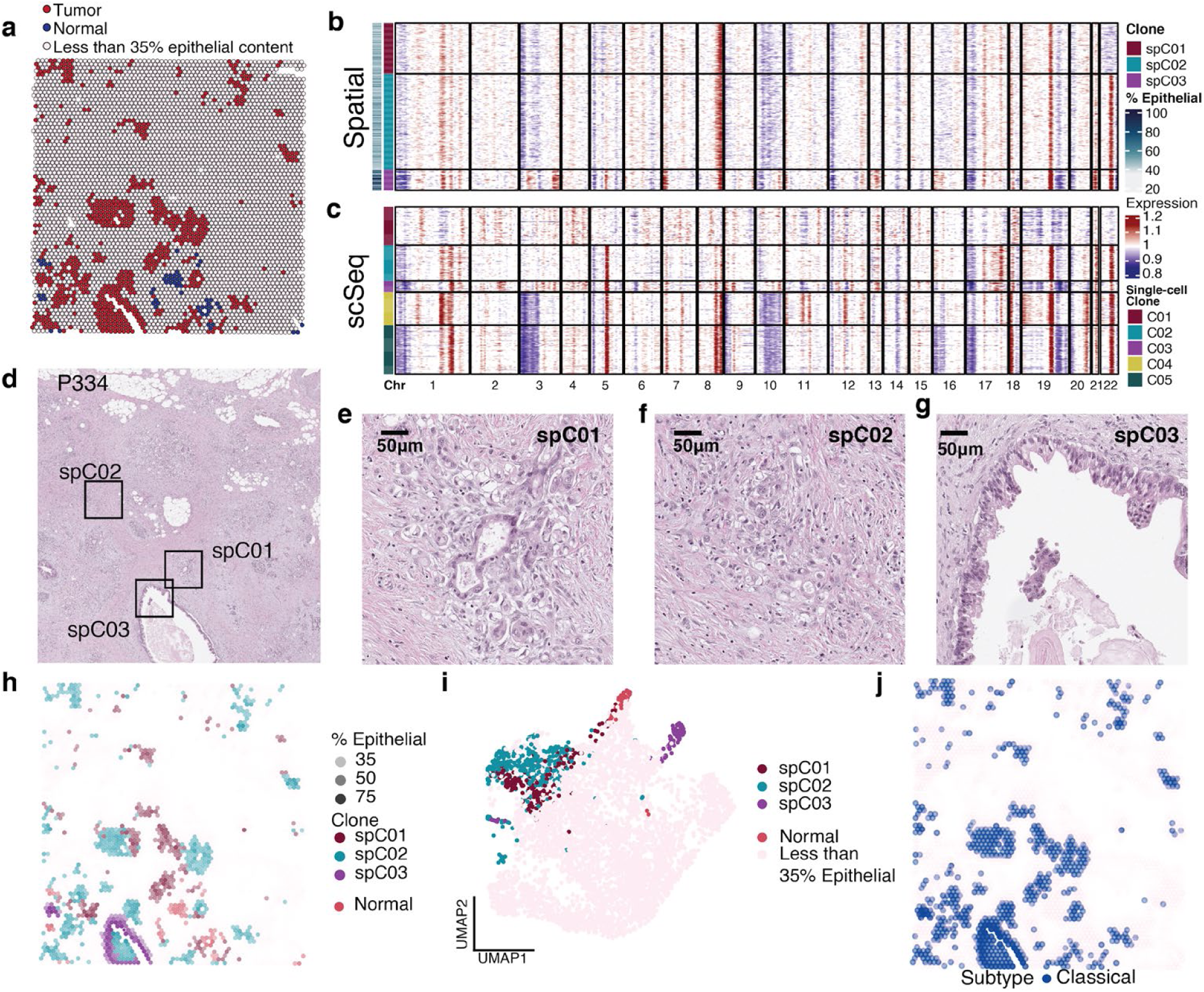
Spatial clonal mapping in PDAC. **a,** Spots with more than 35% epithelial content, as determined by BayesPrism in the profiled region of a representative sample (P334), colored by malignancy, annotated by a pathologist. **b,** CNV profile of spots of P334 with more than 35% epithelial content. Clustered spots are considered clones. **c,** CNV profile of the corresponding single-cell RNA sequencing data from P334. **d,** H&E image of P334 annotated with three clonal regions. **e-g,** H&E images of regions of spCNV1 (e), spCNV2 (f), and spCNV3(g) clones. **h,** Spatial mapping of the clones in the profiled region of P334. **i,** UMAP of the spots in the profiled region of P334, color-coded by clone. **j,** Spatial mapping of the subtypes of the clones of P334.

In a representative example (P334), we identified three distinct clones in the spatial data, each exhibiting unique characteristics (Fig. 6b,d). Clones spC01 and spC02 were more intermixed with stromal cells (Fig. 6e,f,h), while clone spC03 appeared to have differentiated into a more ductular/glandular histology (Fig. 6g,h). When projected onto the UMAP of the sample, spC03 clustered further from the other two clones, likely due to its stronger epithelial signal, as its spots contained higher epithelial content (Fig. 6i). Finally, in this specific example, all clones were of the classical subtype (Fig. 6j), consistent with the matched scRNA-seq analysis.

### Dispersed clonal populations drive lymph node invasion and shift toward basal phenotype in PDAC

Across all samples, we observed two distinct patterns of spatial clone distribution: some clones formed localized clonal neighborhoods, while others were dispersed throughout the tissue (Fig. 7a, Suppl. Fig. 6a). To quantify this phenomenon, we applied local Moran’s I (LMI), a spatial statistic that evaluates the likelihood of a variable being randomly distributed in space. An LMI value near zero indicates random distribution, a significant positive LMI suggests localization, and a significant negative LMI indicates dispersion. Using this metric, we identified clones that were predominantly localized (Fig. 7b, Suppl. Fig. 6b) as well as clones that exhibited both localized and dispersed distributions in different regions of the tissue (Fig. 7c, Suppl. Fig. 6c).

**Figure 7.**
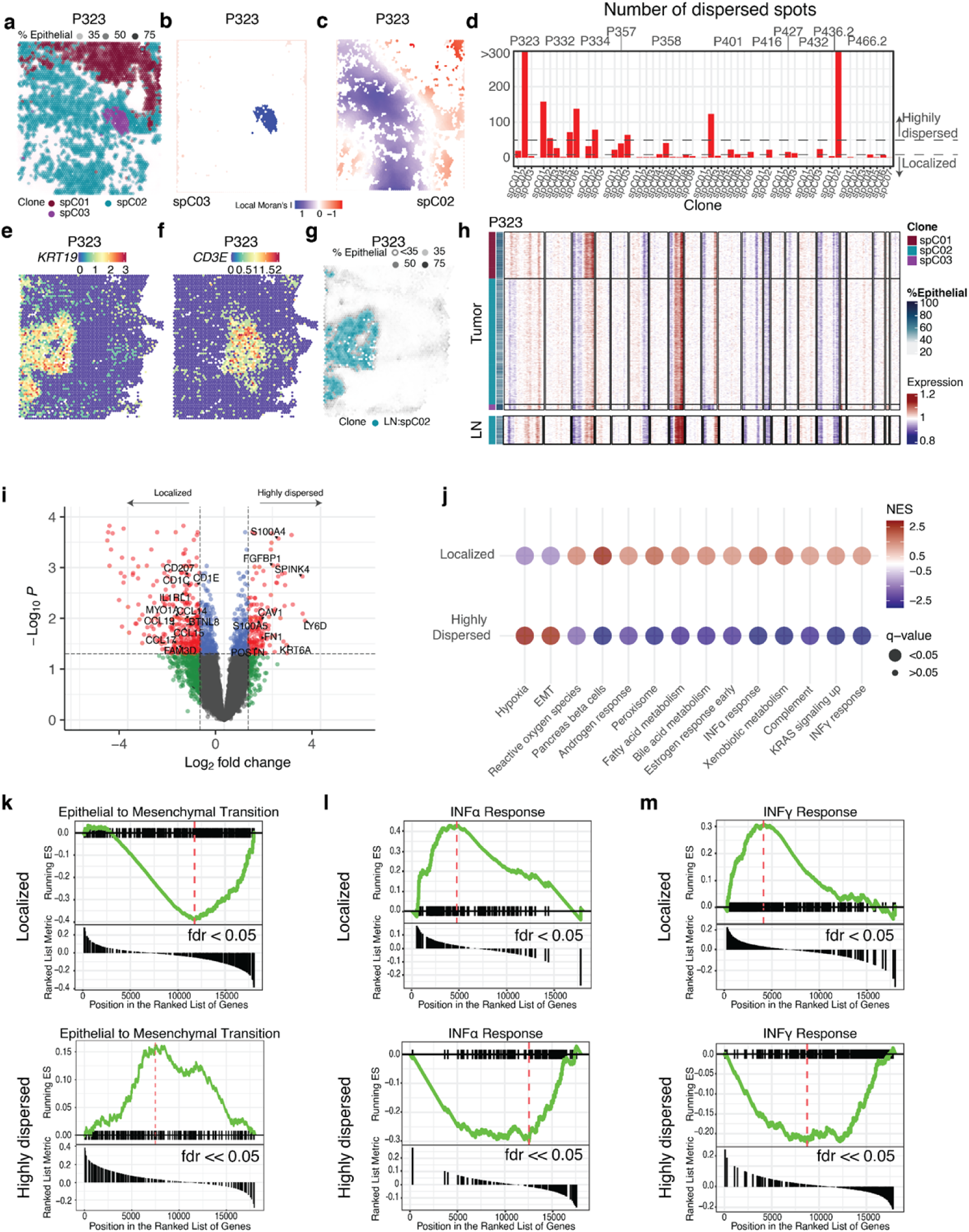
Dispersed clonal populations facilitate lymph node invasion in PDAC. **a,** Spatial mapping of the clones in the profiled region of a representative sample (P323). **b,** Local Moran’s I score for clone spCNV2 of P323. **c,** Local Moran’s I score for clone spCNV3 of P323. **d,** Barplot showing the number of dispersed spots for each clone of each sample. **e,** *KRT19* expression in the lymph node of P323. **f,** *CD3E* expression in the lymph node of P323. **g,** Spatial mapping of spCNV2 clone that has invaded P323’s lymph node. **h,** CNV profile of spots of the tumor region (top) and the lymph node (bottom) of P323 with more than 35% epithelial content. Clustered spots are considered clones. **i,** Volcano plot of differentially expressed genes in highly dispersed and localized clones. **j,** Top 15 pathways enriched in localized and highly dispersed clones. **k,** Enrichment plot of epithelial to mesenchymal transition hallmark pathway for localized (top) and highly dispersed clones (bottom). **l,** Enrichment plot of Interferon alpha response hallmark pathway for localized (top) and highly dispersed clones (bottom). **m,** Enrichment plot of Interferon gamma response hallmark pathway for localized (top) and highly dispersed clones (bottom).

The application of the LMI statistic revealed a mixture of dispersed and localized clones across all samples (Fig. 7d, Suppl. Fig. 5d). To further categorize these clones, we classified them based on the number of dispersed spots they inhabited: clones with more than 50 dispersed spots were classified as highly dispersed, those with fewer than 10 dispersed spots as localized, and clones with 10–50 dispersed spots as having low dispersion. This classification identified 9 of 51 clones as highly dispersed across 7 samples, 17 clones with low dispersion, and the majority, 25 clones, as localized (Fig. 7d).

We hypothesized that dispersed clones represent those with higher metastatic potential. To test this hypothesis, we further profiled two samples that contained primary tumor and regional lymph node metastases (P323, P334). In both cases, we identified a single clone that had metastasized from the primary tumor to its adjacent lymph node (Fig. 7e-g, Suppl. Fig. 6e-g). Notably, this is the same clone that exhibited a highly dispersed spatial distribution within the primary tumor (Fig. 7h, Suppl. Fig. 6h).

Given that dispersed clones have a higher propensity to metastasize, and considering the established link between the basal subtype and increased EMT activity and metastasis, we sought to investigate the subtypes of clones in our spatial dataset. Across the dataset, most clones were classified as predominantly classical, including all nine highly dispersed clones. Notably, six out of nine highly dispersed clones, including both clones confirmed to have metastasized to the lymph node, exhibited amplification of the *MYC* locus (Suppl. Fig. 5).

Building on these observations and our earlier finding that the classical-to-basal transition is accompanied by immune evasion, we hypothesized that dispersed clones may represent a transitional state, wherein some cells are shifting toward a basal-like phenotype. To further test this hypothesis, we assessed the transcriptional and functional changes associated with dispersed clones, focusing on EMT activity and the depletion of immune-related pathways.

DGEA analysis of the localized and highly dispersed clones, revealed that markers of the basal signature (*LY6D, KRT6A, KRT15, FGFBP1*) and the EMT signature (*FN1, S100A4, S100A5*) were upregulated in the dispersed clones (Fig. 7i, Suppl. Fig. 6i), despite their overall classical character. In contrast, immune-related genes (*CD207, CD1C, CD1E, CCL15, CCL17, CCL19*) were overexpressed in the localized clones, suggesting that cells within the dispersed clones are transitioning toward a more basal phenotype (Fig. 7i, Suppl. Fig. 6i).

Given that spatial transcriptomics spots typically contain more than one cell and considering the high abundance of myCAFs in the tissue, we tested for differences in cell type composition between the highly dispersed and localized clones. This analysis revealed a comparable proportion of both epithelial cells and myCAFs in the two groups (Suppl. Fig. 6j), supporting the notion that the observed markers are primarily driven by the epithelial cells within the clones rather than other cell types.

Finally, GSEA revealed that dispersed clones were enriched for pathways associated with EMT and hypoxia, suggesting an enhanced capacity for invasion, adaptation, and metastasis (Fig. 7j,k). In contrast, localized clones demonstrated enrichment for immune-related pathways such as interferon alpha and gamma responses (Fig. 7j,l,m). These findings indicate that localized clones are subject to stronger immune surveillance, while dispersed clones leverage mechanisms such as EMT to facilitate dissemination and immune evasion. Together, these results support the notion that dispersed clones may represent a transitional state, progressively shifting toward a more basal and aggressive phenotype characterized by enhanced EMT and immune evasion.

## DISCUSSION

The comprehensive single-cell and spatial transcriptomics analyses of PDAC presented in this study highlight the intricate interplay between clonal evolution, transcriptional phenotypes, and TME interactions. By analyzing a unique cohort of 62 primary PDAC tumor samples, the largest published to date, we uncover key mechanisms driving tumor heterogeneity, including clonal dynamics and spatial distribution patterns, offering new insights into the biology of this highly aggressive cancer.

Our findings highlight the substantial heterogeneity within and across PDAC tumors. The identification of multiple clones within individual patient samples reveals diverse genetic and transcriptional landscapes, shaped by distinct CNV profiles, consistent with previous reports^20,46^. These results reinforce the dynamic nature of clonal evolution, demonstrating how genomic alterations contribute to the development of diverse transcriptional states within tumors. This heterogeneity is further reflected in the coexistence of basal and classical transcriptional subtypes within individual tumors. The observation of a transcriptional continuum, rather than a strict binary classification, aligns with the prior work of our group and others^18,21,47,48^ and underscores the complex architecture of PDAC tumors.

Clonal heterogeneity also influenced interactions with the TME, as demonstrated by the LRI analysis. Distinct clones exhibited unique interaction patterns with immune and stromal cell populations, underscoring the role of clonal diversity in shaping the tumor’s immunosuppressive microenvironment. Notably, CNV-driven alterations played a critical role in modulating these interactions, with amplifications and deletions affecting key ligand-receptor pairs. For example, amplification of PVR and NECTIN2 enhanced immune checkpoint interactions with TIGIT, potentially contributing to immune evasion. In contrast, deletions of the 9p24 locus, that includes PDCD1LG2 and CD274, led to the suppression of PD-1/PD-L1 interactions, suggesting a diminished role for this checkpoint axis in PDAC. These CNV-driven alterations in cell-cell interactions highlight the need for therapeutic strategies tailored to the genetic landscape of individual tumors.

Analysis of matched pre- and post-treatment samples revealed that chemotherapy further influenced clonal dynamics, driving distinct patterns of expansion and elimination. Clones that persisted post-treatment exhibited increased interactions with iCAFs and endothelial cells, suggesting that these supportive niches contribute to their survival. These findings underscore the critical role of the TME in shaping clonal behavior and highlight the potential of targeting these interactions to enhance therapeutic efficacy. While the limited size of our matched cohort calls for further investigation to fully characterize these processes, it nonetheless provides valuable insights into the molecular and interactional dynamics of clonal survival.

A key finding of this study is the identification of an amplification of 8q24, the locus that includes the MYC oncogene, as a hallmark of classical-to-basal transitions. This transition was accompanied by a downregulation of immune-related pathways, indicating a shift in the tumor’s immune landscape that likely facilitates immune evasion and contributes to therapy resistance. Functional experiments further supported this observation, showing that Myc knockdown in basal murine PDAC cell lines shifted their transcriptional phenotype toward a classical signature while enhancing immune response pathways. These results underscore the role of *MYC* and immune regulation in promoting cellular plasticity and facilitating the transition to a more aggressive basal phenotype. These transitions are likely critical for disease progression, as basal tumors are associated with worse clinical outcomes. Given *MYC*’s well-established role as an oncogene, these findings reinforce its potential as a promising target for therapeutic intervention in PDAC.

LRI analysis in transitioning pairs of classical and basal clones further revealed that classical clones, prior to transitioning, interacted with *SPP1*+ TAMs through the APLP2-PIGR pathway. The downregulation of this pathway during the transition may facilitate reduced immune surveillance, promoting a permissive tumor microenvironment for the emergence of basal clones. A recent study suggests that PIGR enables the transcytosis of dimeric IgA across tumor epithelial cell surfaces, which can neutralize intracellular oncogenic drivers like KRAS^G12D^ and expel them from the cel_l_^40^. This highlights PIGR’s potential role in modulating tumor-immune interactions and suggests it could be explored as a therapeutic target in epithelial-derived classical tumors. In contrast, basal clones that emerged from these transitions established interactions with TAMs and stromal cells via pathways such as SEMA3C-NRP2, TGFA-EGFR, and GAS6-AXL. These pathways are associated with enhanced tumor progression, immune evasion, and a more invasive phenotype, emphasizing their therapeutic potential. Moreover, the involvement of receptors such as NRP2, and AXL in these interactions provides additional therapeutic avenues. NRP2, a co-receptor implicated in angiogenesis and lymphangiogenesis, has been linked to cancer progression and metastasis^41,42^, positioning it as a promising candidate for drug development. AXL, a receptor tyrosine kinase, is associated with EMT, immune evasion, and drug resistance, with inhibitors such as bemcentinib (BGB324) and dubermatinib, undergoing clinical evaluation for multiple cancers, including pancreatic cancer^49^.

We also profiled 13 samples, including 11 primary PDAC tumor samples, and 2 lymph nodes with metastasis, with spatial transcriptomics, all matched with the single-cell data. While previous studies have identified clones using either single-cell or spatial transcriptomics^20,22,50^, our study identified the same CNVs in both modalities. Moreover, we successfully identified identical clones in a subset of samples. The inability to identify identical clones in all cases may be attributed to selection bias, such as differences in sampling regions or processing techniques.

The spatial analysis of PDAC tumors revealed distinct localized and dispersed patterns of clone distribution in space. Dispersed clones showed a strong association with metastatic potential, as evidenced by their presence in both primary tumors with and matched lymph node metastases. These clones exhibited elevated EMT activity, reinforcing their role in tumor dissemination. The role of EMT in promoting metastasis has been widely documented, with evidence linking EMT activation to transcription factors such as ZEB1 and SNAIL as well as the MYC oncogene^51–54^. MYC amplification, identified in our study as a hallmark of classical-to-basal transitions in our study, was present in six out of nine dispersed clones. Moreover, EMT activation has been associated with immune evasion and resistance to therapy^48^, emphasizing the significance of the dispersed clones in shaping disease progression and treatment outcomes. To investigate further, we analyzed the genes and pathways differentially expressed between localized and dispersed clones. Although dispersed clones were predominantly classical, they showed enrichment for basal markers and pathways associated with cellular plasticity. In contrast, localized clones were enriched in immune-related pathways. These findings suggest that dispersed clones may represent a transitional state toward a more aggressive basal phenotype, contributing to metastasis and tumor progression.

Overall, our findings provide a much more comprehensive view of the molecular and spatial heterogeneity in PDAC than currently exists, highlighting the critical role of clonal diversity in shaping tumor progression. Mechanisms such as classical-to-basal transitions driven by MYC amplification, the immune evasion strategies employed during these transitions, and the metastatic potential of dispersed clones emphasize the complex interplay between clonal evolution, transcriptional plasticity, and the TME. The distinct interaction patterns between clones and their microenvironment further underscore the challenges in targeting this heterogeneity therapeutically in PDAC patients. By identifying key pathways and genetic alterations driving these processes, our study offers valuable insights that could inform the development of more effective therapeutic strategies to combat this aggressive disease.

## METHODS

### Sample collection and preparation

In total, 59 patients were recruited between August 2020 and April 2023 at the Perlmutter Cancer Center at New York University (NYU) Langone Health as part of a study approved by the NYU Langone Health Institutional Review Board. Patients gave written informed consent for sample collection and sequencing, as well as clinicopathological data collection. This cohort is composed of 62 samples including 24 samples collected via surgical resection and 38 via biopsy. Clinical somatic tumor analyses were performed using the Tempus xT596 gene panel, Tempus Labs, Chicago, IL and FoundationOne CDx 324 gene panel, Foundation Medicine, Cambridge, MA.

### Single-cell library preparation, sequencing, and data processing

Samples were processed to isolate single cells as previously described by our group^18^ before the single cell suspensions were processed by the Genome Technology Center at NYU School of Medicine using 10x Genomics Chromium single-cell 3’ Reagent Kits v3, or Chromium single-cell 5’ Reagent Kits v2 following manufacturer guidelines (samples sequenced after August 2021), and run on the Illumina NovaSeq 6000 system (Illumina, San Diego, CA). Following conversion of output BCL files via Illumina’s bcl2fastq, the FastQ files were processed with 10x Genomics Cell Ranger v7.0.1^55^ and aligned to the reference hg38/GRCh38 reference genome. Non-PCR duplicated reads that confidently mapped to the reference genome with valid unique molecular identifiers (UMIs) and barcodes were used to generate a counts matrix. Raw data were initially filtered to remove ambient RNA using SoupX v1.5.258. Seurat v5^56^ and scooter^57^ packages were used to perform downstream data analysis, including the identification of highly variable genes, dimensionality reduction, standard unsupervised clustering algorithms, visualization, and the discovery of differentially expressed genes. Cells with >1200 UMIs, >500 genes, and <15% mitochondrial genes, were retained. Counts data was normalized by total gene expression followed by log-transformation with a scaling factor of 10,000. scDblFinder v1.6.0^58^ was used to identify and remove putative doublets and multiplets present in the data. Clusters were calculated using the first 30 PC dimensions and 30 nearest neighbors and visualized with UMAP projection.

The dimensionality of the scaled non-batch corrected data matrix of the combined data of the samples was reduced to project the cells in two-dimensional space, using principal component analysis followed by UMAP. A combination of different clustering resolutions (0.1 to 1.9) and marker gene expression were used to identify cells grouped together by classic marker genes. Markers used for both broad and granular cell types (Ext. Fig. 1a,b) were the same as those used in previous reports from our group^18^. The rest of the analysis was performed on data from each individual sample, using the cell type annotation from the combined data matrix.

### Identification of clones and malignant epithelial cells

Clone identification was performed for each of the samples independently with inferCNV^28^. More specifically, all epithelial cells in each sample were tested for CNVs (observation set), with 90% of non-epithelial cells used as the reference normal set, and the remaining 10% of non-epithelial cells included in the observation set, as an internal control. To ensure robustness, we focused on samples containing more than 100 epithelial cells, resulting in a final study cohort of 50 samples. An exception to this criterion was made for matched pairs of samples collected before and after chemotherapy, where combining these pairs provided enough cells for analysis. InferCNV uses a Hidden Markov Model (HMM) and Bayesian model to calculate the probability of an identified CNV to be real. From the identified CNVs, we kept all the CNVs that are shown to be real with a probability higher than 0.7. Hierarchical clustering of the CNV profile of each cell revealed clusters of cells that shared the same CNVs, that were considered clones. In order to distinguish malignant cells from normal epithelial cells, we assigned a score on each clone based on the percentage of each chromosome that is part of a CNV. We then compared the score of each epithelial cell to that of the control non-epithelial cells; epithelial cells with a score higher were considered malignant. For each clone, CNVs that were called from inferCNV were mapped to the genomic positions of cytogenetic bands.

### Spatial transcriptomics library preparation, sequencing, and data processing

Samples were obtained via surgical resection and confirmed by a pathologist to contain sufficient tumor content. Spatial transcriptomics profiling was performed using the 10x Genomics Visium CytAssist platform on FFPE samples, following the manufacturer’s protocol. After tissue sectioning and placement on Visium slides, spatially barcoded cDNA libraries were generated using 10x Genomics Visium for FFPE Library Preparation Kits. Libraries were sequenced on the Illumina NovaSeq 6000 system.

Raw sequencing data were processed with 10x Genomics Space Ranger v2.1.1, aligning reads to the GRCh38/hg38 human reference genome. Spatial gene expression matrices were generated by mapping transcripts to their corresponding tissue locations. Deconvolution was performed using the R package BayesPrism^59^, with scRNA-seq data serving as the reference for cell type identification. Spots were filtered to include only those with >1000 UMIs, >500 detected genes, and <10% mitochondrial gene content. Downstream processing, including normalization, dimensionality reduction, clustering, and visualization, was performed using Seurat v5.

### Clone identification in spatial transcriptomics data

Clone identification in spatial transcriptomics data was performed using inferCNV. In spatial data, spots containing more than 35% epithelial cells (as determined by BayesPrism deconvolution) were used as the observation set, while spots with less than 10% epithelial content served as the reference data. All subsequent steps for clonal identification were identical to those applied in the scRNA-seq analysis.

### Whole genome sequencing and analysis

In cases where fresh frozen tissue was available, the inferred CNVs were further validated with low passage WGS (median read depth 1x). FastQ files were processed using the Seq-N-Slide analysis workflow for WES/WGS^60^. Briefly, FastQ files were trimmed with Trimmomatic v0.36 (trailing=5, sliding window=4:15, minlen=35). Next the reads were aligned to the reference hg38 genome with BWA-MEM alignment v0.7.17, filtering for a mapping quality of at least 10. Duplicates were removed by sambamba v0.6.8. Filtered bam files were then realigned and recalibrated with GATK v3.8.1. CNVs were predicted by Control-FREEC v11.6^61^ using default parameters for WGS. From this, the BedGraph formatted tracks of CNVs were plotted in a heatmap, aligned to the inferCNV profile.

### Cell type composition analysis

To compare differences in cell type composition across conditions, we used the propeller^62^ R package, which employs an empirical Bayesian framework to stabilize variance estimates when analyzing compositional data. Statistical significance was determined using FDR correction for multiple comparisons.

### Identification of subtypes

Individual cells were profiled using published subtype gene signatures^12–15^ to generate a module enrichment score for each subtype for each cel_l_^56^. Pearson correlation analysis revealed that gene expression signatures corresponding to the basal and classical subtypes were highly correlated with previously published subtype-specific signatures. Specifically, basal subtype signatures showed strong correlations with other basal-equivalent signatures, and classical subtype signatures exhibited high correlations with classical-equivalent signatures. Based on this, we selected only the Moffitt study’s basal and classical subtype signatures for downstream analysis^12^. To classify individual cells, per-cell basal and classical signature scores were compared. If at least one of the scores was positive, the cell was assigned to the subtype with the higher score. Conversely, if both scores were negative, the cell was classified as expressing “neither” subtype signature.

To classify clones by subtype, we applied a majority-rule approach based on the predominant classification of individual cells within each clone. A clone was assigned to a major subtype (basal or classical) if at least 50% of its cells belonged to that subtype and the difference in prevalence between the two subtypes exceeded 7.5%. If cells expressing neither the classical nor the basal subtype accounted for at least 50% of the clone, the clone was classified as having no major subtype. Finally, clones where the proportion of both major subtypes fell between 33% and 50% - without either reaching the 50% threshold- or the difference between the two subtypes was less than 7.5% were classified as “Mixed”.

### Pathway Analysis

Pathway analysis was performed individually for each sample with singleSeqGset^63^, using the Hallmark^64^, KEGG^65^, and Reactome^66^ gene set collections, to conduct differential GSEA by comparing each clone against all others within the same sample. This approach identified statistically significant enriched gene sets for each clone. To define recurrently enriched pathways, gene sets enriched in more than 5% of clones were selected. Pearson correlation analysis was then used to assess similarity among gene sets, clustering them into broader pathway groups based on their correlation structure. Each clone was subsequently assigned to a dominant pathway group, determined by the most enriched pathway group in that clone.

### Phylogenetic tree construction

A phylogenetic tree was constructed for each of the tumors, based on the CNVs from the scRNA-seq and the spatial transcriptomics data generated by inferCNV^29^. The phylogenetic trees were built manually, by identifying CNVs that are shared across clones, assuming that a copy number alteration is irreversible once occurred. Normal, non-malignant epithelial cells served as the root of the phylogenetic trees and identified clones occupying both nodes. Nodes are scaled in size proportional to the number of cells present in that specific clone. Shared CNVs among clones served as evidence of a common ancestry between clones, informing the tree-building process. In cases where two clones shared CNVs and both had additional CNVs not detected in the other one, we assumed that these clones shared an unobserved common ancestor from which these clones evolved due to the newly gained CNV events. Unobserved clones are set to a standardized node size due to no cells observed in that node. Phylogenetic tree branches are solid lines between clones that are directly observed and dashed to and from unobserved nodes given their inferred status. Distance between nodes is proportional to the clone score representing the CNV burden on the genome to provide a pseudotime representation of mutational burden. The distance of any node back to the root is equal to that node’s mutational burden score. Therefore, the closer two nodes are, the fewer different CNVs between them, and therefore a lower CNV burden pseudotime.

To visualize all samples in the study at once in the “phylogenetic forest”, plots are simplified to represent all nodes at the same size and with solid branches that do not represent the clones’ mutation scores. Nodes are still representative of the Moffitt subtype of the represented clone.

### Ligand-receptor interaction analysis

To assess the influence of malignant clonal evolution on the TME in PDAC, we performed LRI analysis for each individual sample, analyzing interactions between clones and TME cell types using CellPhoneDB v5.0.0^67^. A minimum of 10 cells per cell type per sample was required for inclusion in the analysis.

To evaluate how CNVs influence ligand-receptor interactions, we first mapped ligands and receptors from the CellPhoneDB database to their respective cytogenetic bands. We then quantified how frequently each gene in an interaction pair was affected by a CNV and how often the corresponding clone-TME interaction was significant. For locus amplifications, we calculated an enrichment score as the product of the number of times the gene was amplified and the number of times the interaction in which the gene participates was significant. For locus deletions, we calculated a suppression score as the product of the number of times the gene was deleted and the number of times the interaction was not significant.

### Differential Gene Expression and Gene Set Enrichment Analysis after pseudobulking

DGEA was performed on pseudobulk expression data using DESeq2 (reference), in three analyses: classical-to-basal transitions, matched pre- and post-treatment samples, and comparisons between dispersed and localized clones in spatial transcriptomics data. For comparisons involving independent groups, standard DESeq2 workflows were applied, while paired analysis was used for dependent variables, such as the comparison of the clones pre- and post-the classical-to-basal transition, to account for sample-specific effects. GSEA was performed with ClusterProfiler^68^, using the hallmark gene set collection from Molecular Signatures Database (MSigDB_)_^64^ to identify biological pathways enriched in differentially expressed genes.

### Survival analysis

Patients were stratified based on the presence or absence of clones harboring key CNVs identified as prevalent in the transition from classical to basal clones, specifically amplifications of 8q24, 17q21, and 20q13, and deletion of 1p36. After classifying patients according to this genomic signature, survival analysis was performed using the surviva_l_^69^ and survminer^70^ R packages. Kaplan-Meier curves were generated to compare differential survival between groups, and statistical significance was assessed using the log-rank test p-value.

### KMCE murine experiment Cell line

The KMC314J cell line was generated from PDAC of Ptf1a-CreER; KrasG12D; Rosa26MycWT/MycWT mouse and maintained in conditionally reprogrammed culture (CRC) medium (Liu et al, AJP 2012). SMARTvector Inducible Lentiviral shMyc (horizon discovery; V3SM7671-233031217) were transfected to the KMC314J cell line according to manufacturer’s protocol. Briefly, cells were transducted at 0.3TU/MOI in 96 well plate and selected by 2ug/ml of puromycin. Then single cell clones were obtained after drug selection to maximize knockdown efficiency. 2ug/ml of Doxycycline was used to knockdown. Two well knock-down clones were chosen for further experiments by Western blotting of Myc(Y69) antibody (abcam; ab168727) 72h after Doxycycline induction.

### RNA sequencing

The two clones of the KMC314J shMyc cell line were profiled with RNA sequencing, after 72h after Doxyxycline induction. RNA was obtained by RNeasy Mini kit (Qiagen, Germantown, MD) and submitted to Novogene (Sacramento, CA) for library preparation and sequence. Briefly, mRNA was purified from total RNA using poly-T oligo-attached magnetic beads. After fragmentation, the first strand cDNA was synthesized using random hexamer primers followed by the second strand cDNA synthesis. The library was ready after end repair, A-tailing, adapter ligation, size selection, amplification, and purification. The library was checked with Qubit and real-time PCR for quantification and bioanalyzer for size distribution detection. Quantified libraries will be pooled and sequenced on NovaSeq X plus (Illumina), according to effective library concentration and data amount.

### RNA sequencing data processing

Paired-end fastq sequences were trimmed using Trim Galore (v.0.6.3) and default parameters. Pseudoalignment was performed with kallisto (v.0.44.0) using genome assembly GRCm39 and GENCODE (v.M26) annotation; default parameters were used other than the number of threads. The Bioconda package bioconductor-tximport (v.1.12.1) was used to create gene-level counts and abundances (TPMs). Quality checks were assessed with FastQC (v.0.11.8) and MultiQC (v.1.7). Quality checks, read trimming, pseudoalignment and quantitation were performed via a reproducible snakemake pipeline, and all dependencies for these steps were deployed within the anaconda package management system. DEseq2 (v.1.44.0) was used to obtain normalized counts. Any genes with names starting with ‘Rik’ or ‘Gm[0-9]’, in which [0-9] represents a single digit, were filtered out due to the ill-defined ontology of these genes. Also, genes with unnormalized count>=10 at least 4 out of 16 samples (number of replicates) were included.

### Gene Set Enrichment Analysis

GSEA was performed using the mm10 orthologs of the marker genes of published subtype signatures^12–14,48^ and the Hallmark gene set collection from MSigDB^64^, with gene set permutation and classic enrichment statistic.

## Data availability

scRNA-seq and spatial transcriptomics data is currently being uploaded to dbGaP.

## Acknowledgments

We would like to thank the members of the Center for Biospecimen Research and Development at NYU Langone Health [RRID:SCR_017930] for support with tissue acquisition, the Experimental Pathology Research Lab [RRID:SCR_017928] at NYU Langone Health for the spatial transcriptomics experiments, the Genome Technology Center [RRID:SCR_017929] at NYU Langone Health for expert library preparation and sequencing, and the Applied Bioinformatics Laboratories (ABL) [RRID:SCR_019178] at NYU Langone Health for providing bioinformatics support and helping with the analysis and interpretation of the data. These shared resources are partially supported by the Cancer Center Support Grant P30CA016087 at the Laura and Isaac Perlmutter Cancer Center. This work has used computing resources at the NYU Grossman School of Medicine High Performance Computing (HPC) Facility. This project was supported by NIH R01CA245005 (DMS) and a Sky Foundation seed grant award (DK).

## Author contributions

1. D. K., I.D., A.T. and D.M.S. developed the study concept and were responsible for the study design. D.K. wrote the manuscript, which was then edited by all co-authors. D.M.S., X.J. and C.H. helped acquire human biospecimens. D.L.T., Y.S., D.W., R.H., D.D., P.M., and A.H. processed samples for sc-RNAseq. D.K., D.C.C., E.A.K., and I.D. analyzed the scRNA-seq data. D.K., D.C.C., E.A.K., and I.D. were responsible for the bioinformatics analysis pipelines. X.J. collected patient information and maintained the databases. C.H., D.K., C.L., and D.M.S. analyzed the H&Es. K.R., S.S., C.L., D.D., P.M., and A.H. processed the samples for spatial transcriptomics. D.K. analyzed the spatial transcriptomics data. R.C.S. provided reagents and cell lines for the murine experiment. M.T. and H.Z. conducted the murine experiments. D.K. and M.T. analyzed the data from the murine experiments. T.H.W., C.H., L.W., A.A., M.B., A.W.L. provided study guidance and feedback. T.H.W. and R.C.S. assisted with study design and analysis. D.M.S. and A.T. directed the study. All co-authors approved the final version of the manuscript before submission.

## Competing interests

The authors declare no conflict of interests.

Correspondence and requests for materials should be addressed to D.M.S and A.T..

---

**Extended Figure 1.**
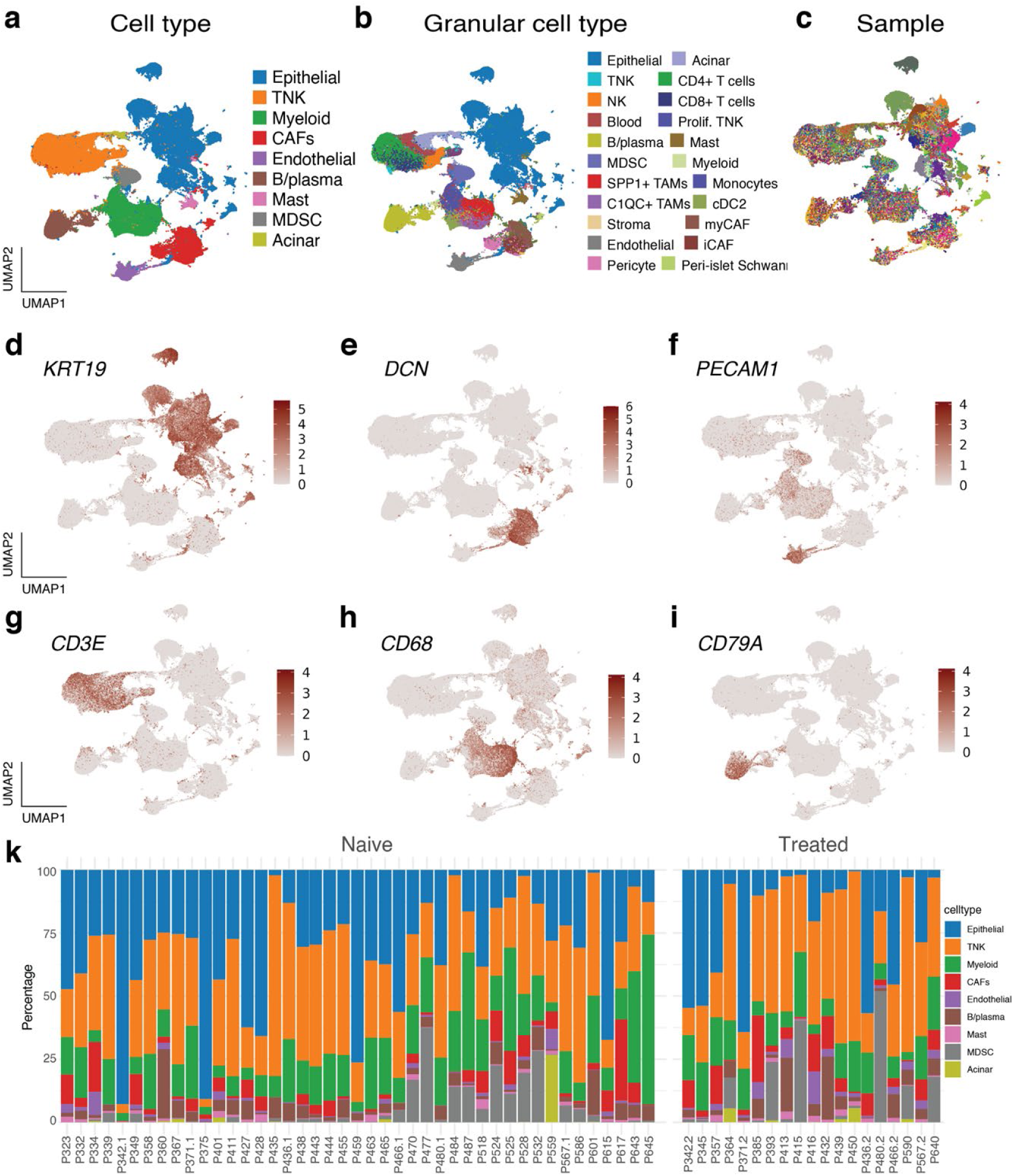
Cell type composition in PDAC. **a,** Non-batch corrected UMAP of all cells, color-coded by major cell type. **b,** Non-batch corrected UMAP of all cells, color-coded by more granular cell type. **c,** Non-batch corrected UMAP of all cells, color-coded by patient, showing patient sample-specific clusters of most epithelial cells and clustering of other cell types across samples. **d-i,** Non-batch corrected UMAP of all cells, showing the expression of the epithelial (*KRT19 - d*), stromal (*DCN -* e), endothelial (*PECAM1 -* f), T (*CD3E -* g), myeloid (*CD68 -* h), and B (*CD79A -* i) cell markers. **k,** Cell type composition per sample.

**Extended Figure 2.**
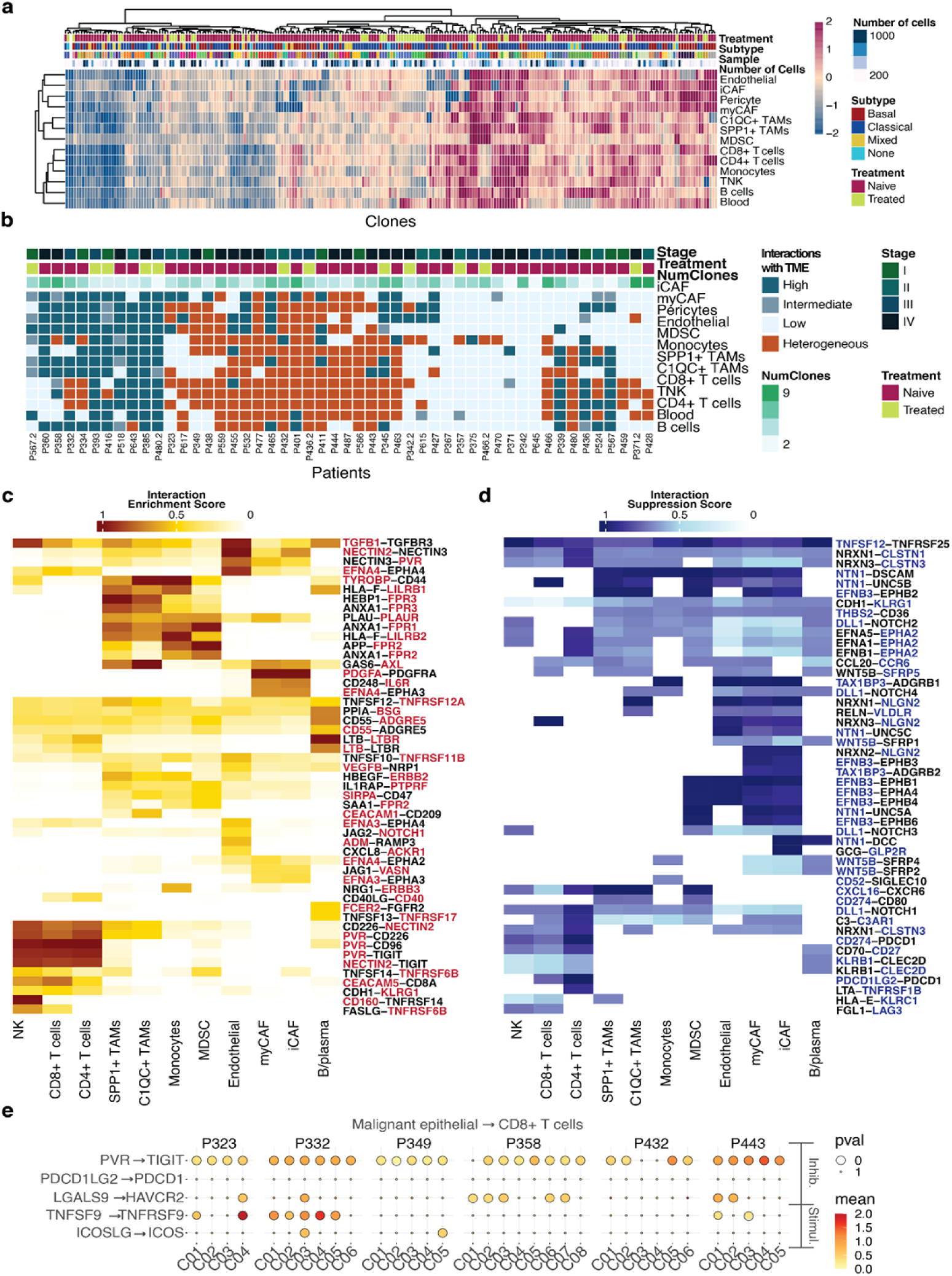
Cell-cell communication analysis reveals clone-specific interactions with other cells in the TME. **a,** Number of interactions between each clone (columns) and other cell types in the TME (rows). **b,** Interaction heterogeneity for each sample. **c,** Heatmap of interactions that were enriched due to an amplification, with red indicating the amplified gene in the interaction. **d,** Heatmap of interactions that were suppressed due to a deletion, with blue indicating the deleted gene in the interaction. **e,** Mean interaction scores for checkpoint molecule LRIs between clones and immune cells of representative samples.

**Extended Figure 3.**
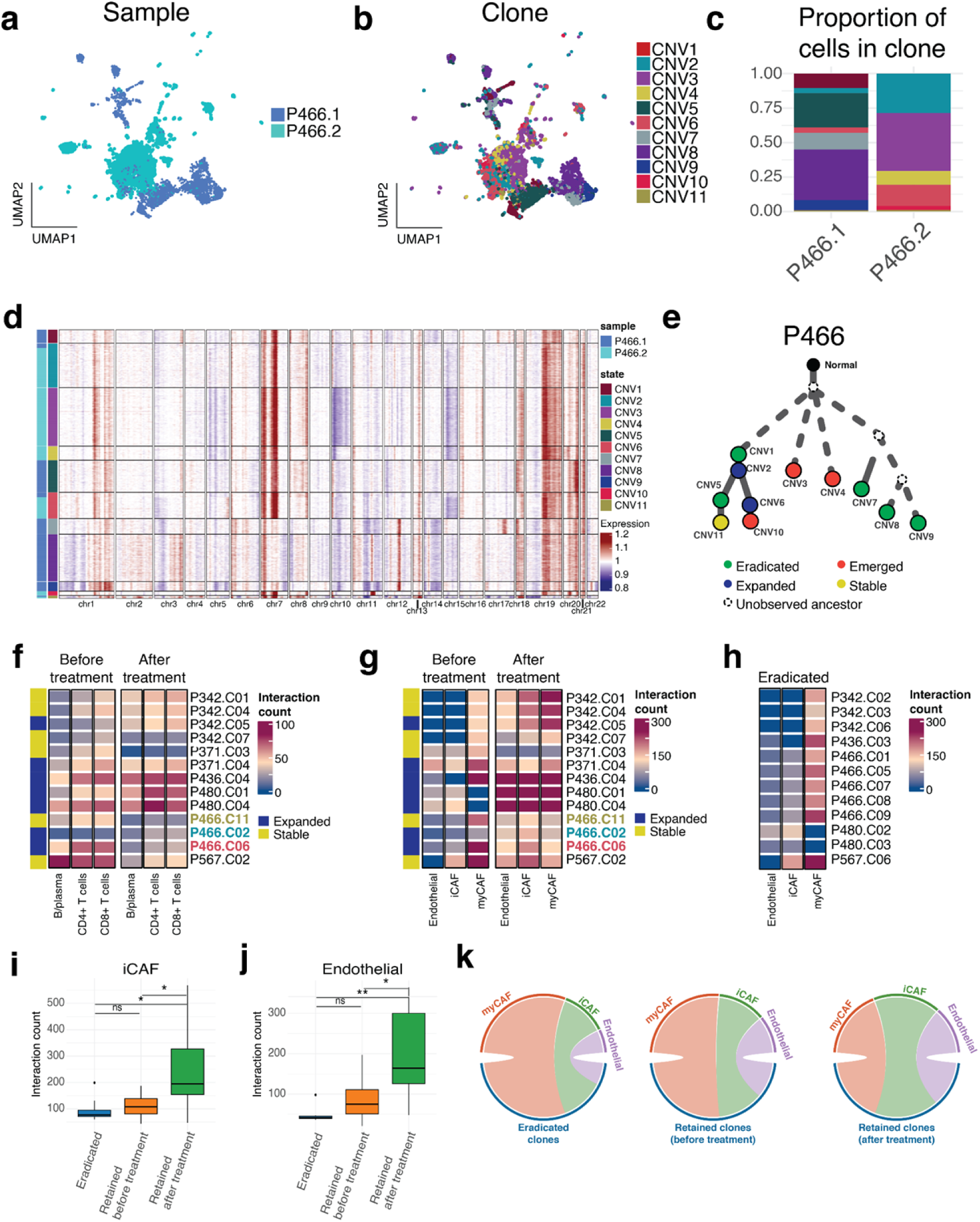
Patterns of clonal dynamics and interactions following chemotherapy. **a,** UMAP showing the malignant epithelial cells of a matched pair of samples collected from the same tumor before (P466.1) and after (P466.2) treatment with chemotherapy, color-coded by sample. **b,** UMAP showing the malignant epithelial cells of a matched pair of samples collected from the same tumor before and after treatment with chemotherapy, colored-coded by clone. **c,** Proportion of clonal cells in each of the matched samples. **d,** heatmap showing the CNV profile of the clones, and how they are distributed in each sample of the matched pair. **e,** Phylogenetic tree of the matched samples shows the patterns of expansion and elimination of clones. **f,** Number of interactions between clones that were retained (either expanded or stable in size) across samples and lymphatic cells. The clones that are common pre- and post-treatment in the example sample are highlighted. **g,** Number of interactions between clones that were retained (either expanded or stable in size) across samples and stromal cells. The clones that are common pre- and post-treatment in the example sample are highlighted. **h,** Number of interactions between clones that eradicated across samples after treatment and stromal cells. **k,** Chord diagrams illustrating interactions between stromal cells and: eradicated clones (left), retained clones before treatment (middle), and retained clones after treatment (right).

**Extended Figure 4:**
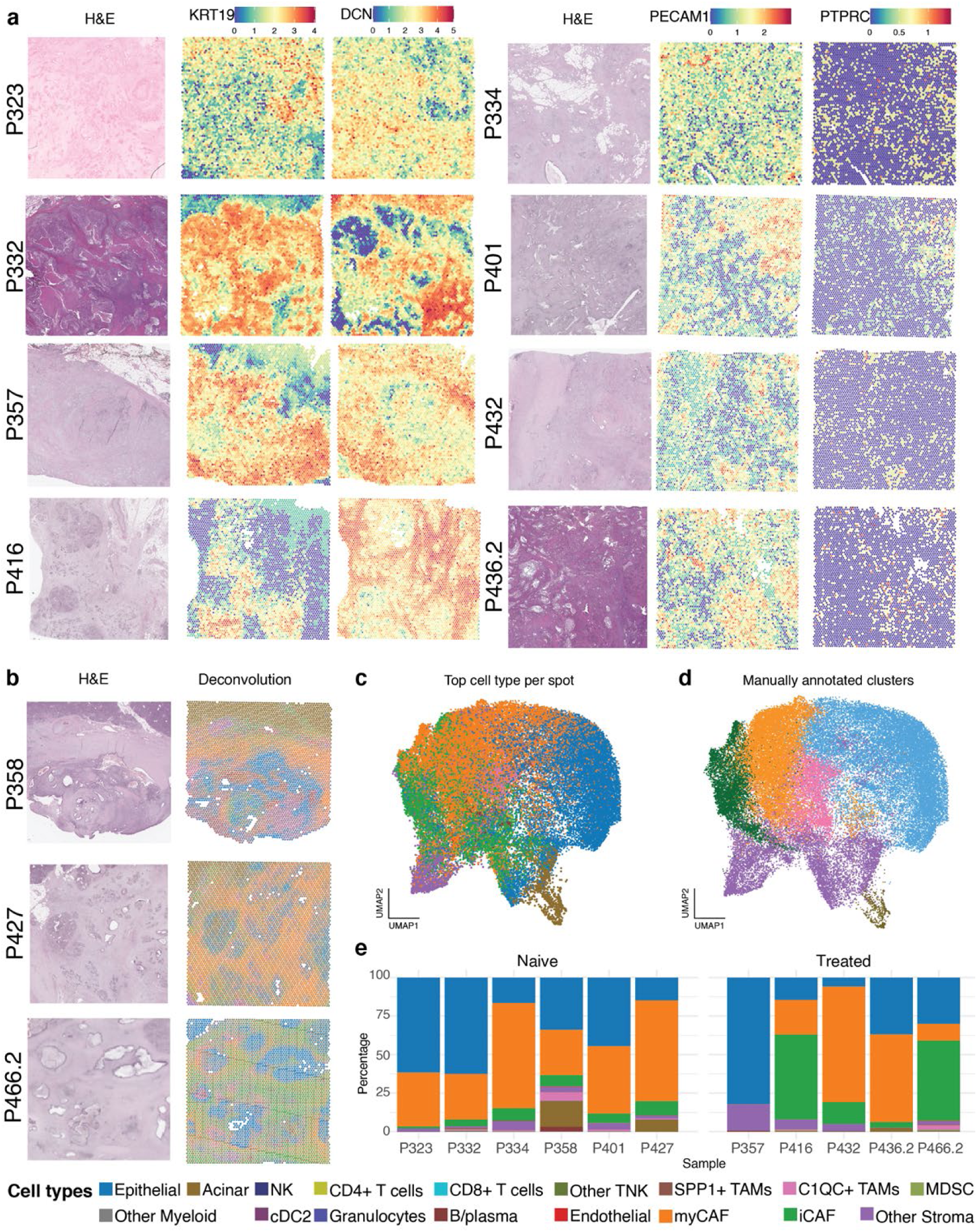
**a,** Histology and common cell type markers for representative samples. **b,** Histology and results of deconvolution for representative samples. **c,** UMAP of all spots, color-coded by the cell type with the highest percentage in each spot, based on deconvolution with BayesPrism. **d,** UMAP of all spots, color-coded by manually annotated clusters. **e,** Top cell type composition per sample.

