## Supplementary material for "Clonal Heterogeneity in Human Pancreatic Ductal Adenocarcinoma and Its Impact on Tumor Progression": Suppl. Fig. 1

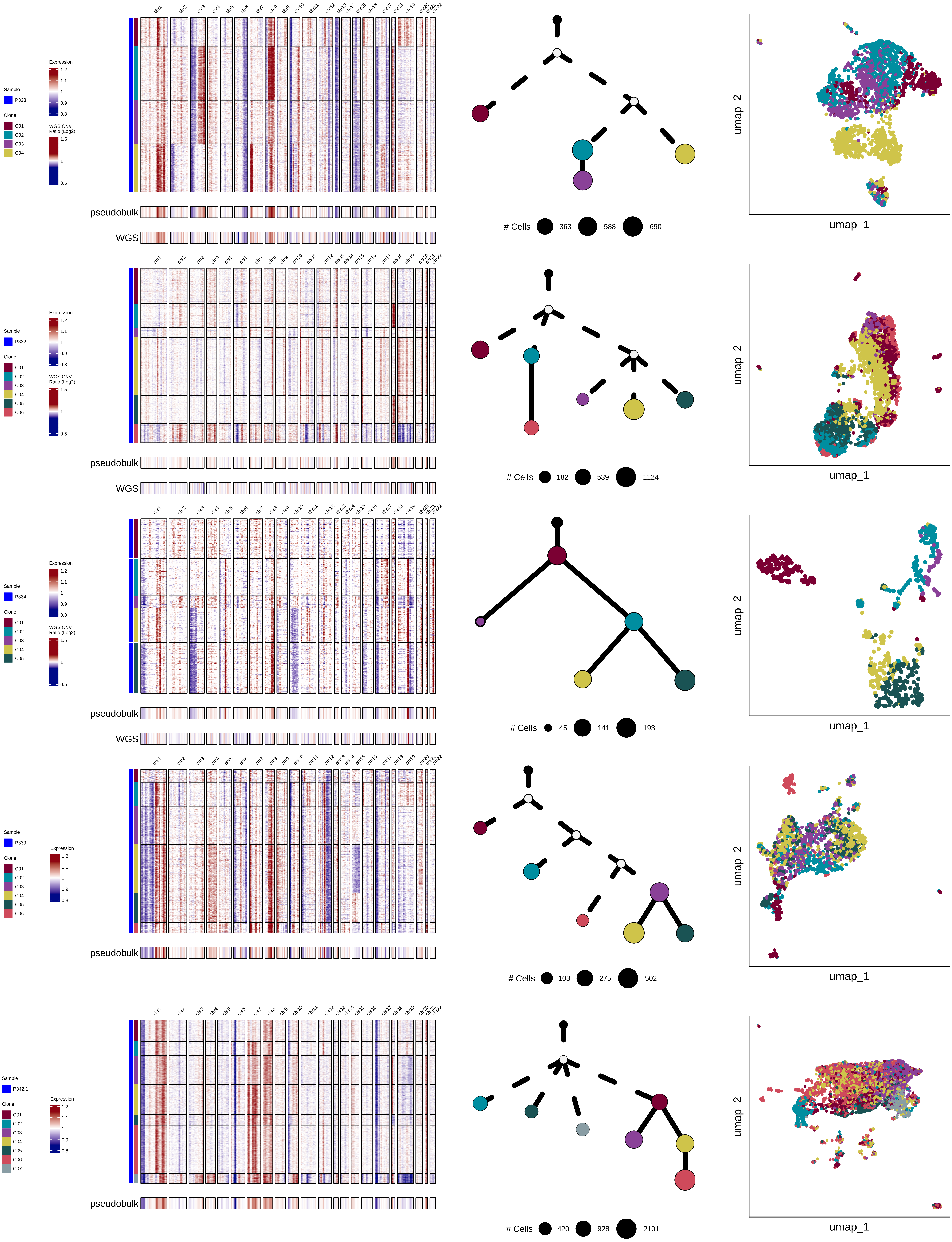

Sample
P323
Clone
C01
C02
C03
C04
Expression
1.2
1.1
1
0.9
0.8
WGS CNV Ratio (Log2)
1.5
1
0.5
chr1
chr2
chr3
chr4
chr5
chr6
chr7
chr8
chr9
chr10
chr11
chr12
chr13
chr14
chr15
chr16
chr17
chr18
chr19
chr20
chr21
chr22
pseudobulk
WGS
### Cells
363
588
690
umap_2
umap_1
Sample
P332
Clone
C01
C02
C03
C04
C05
C06
### Cells
182
539
1124
Sample
P334
Clone
C01
C02
C03
C04
C05
### Cells
45
141
193
Sample
P339
Clone
C01
C02
C03
C04
C05
C06
### Cells
103
275
502
Sample
P342.1
Clone
C01
C02
C03
C04
C05
C06
C07
### Cells
420
928
2101

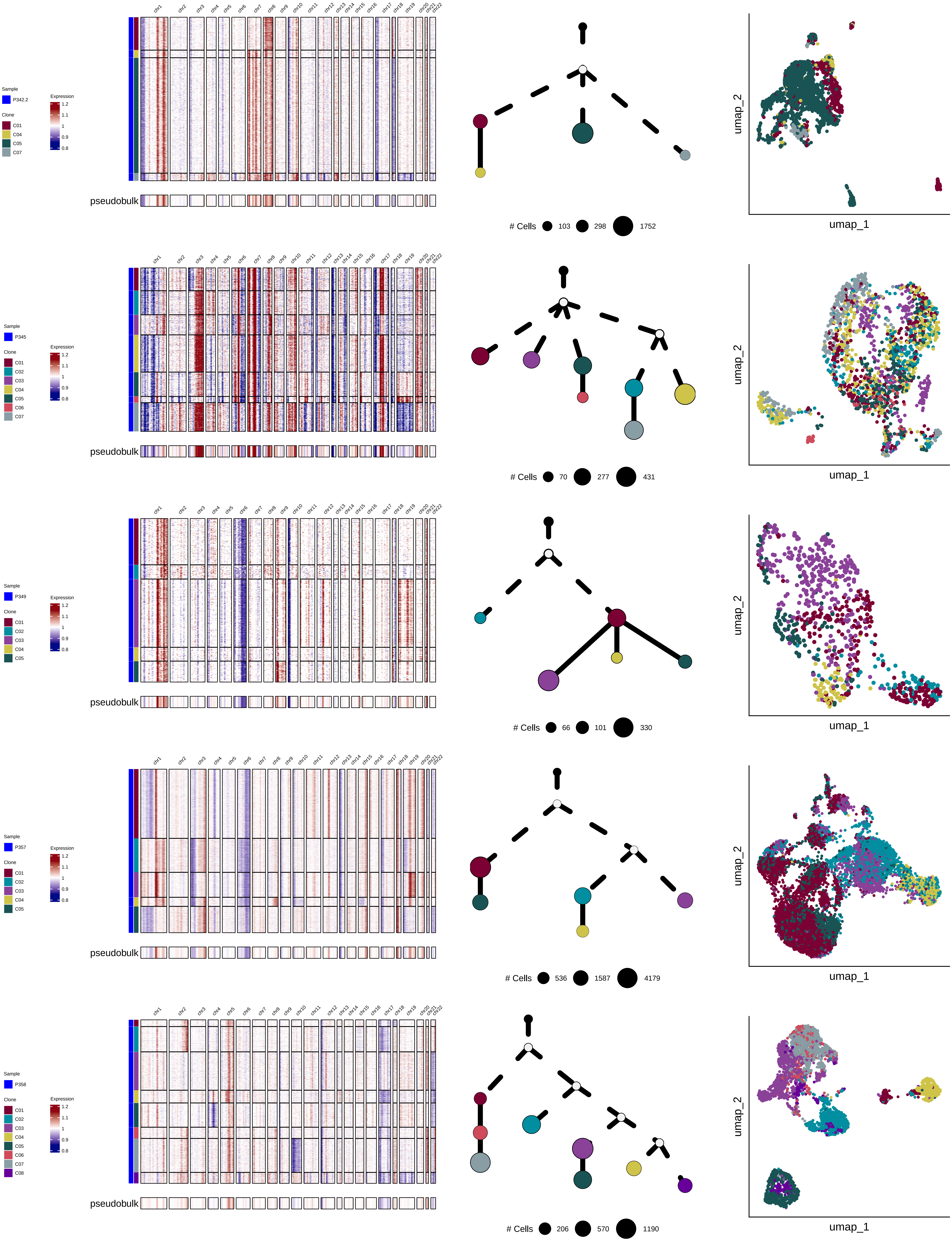
Sample
P342.2
Clone
C01
C04
C05
C07
Expression
1.2
1.1
1
0.9
0.8
chr1 chr2 chr3 chr4 chr5 chr6 chr7 chr8 chr9 chr10 chr11 chr12 chr13 chr14 chr15 chr16 chr17 chr18 chr19 chr20 chr21 chr22
pseudobulk
### Cells 103 298 1752
umap_1
umap_2
Sample
P345
Clone
C01
C02
C03
C04
C05
C06
C07
Expression
1.2
1.1
1
0.9
0.8
pseudobulk
### Cells 70 277 431
umap_1
umap_2
Sample
P349
Clone
C01
C02
C03
C04
C05
Expression
1.2
1.1
1
0.9
0.8
pseudobulk
### Cells 66 101 330
umap_1
umap_2
Sample
P357
Clone
C01
C02
C03
C04
C05
Expression
1.2
1.1
1
0.9
0.8
pseudobulk
### Cells 536 1587 4179
umap_1
umap_2
Sample
P358
Clone
C01
C02
C03
C04
C05
C06
C07
C08
Expression
1.2
1.1
1
0.9
0.8
pseudobulk
### Cells 206 570 1190
umap_1
umap_2

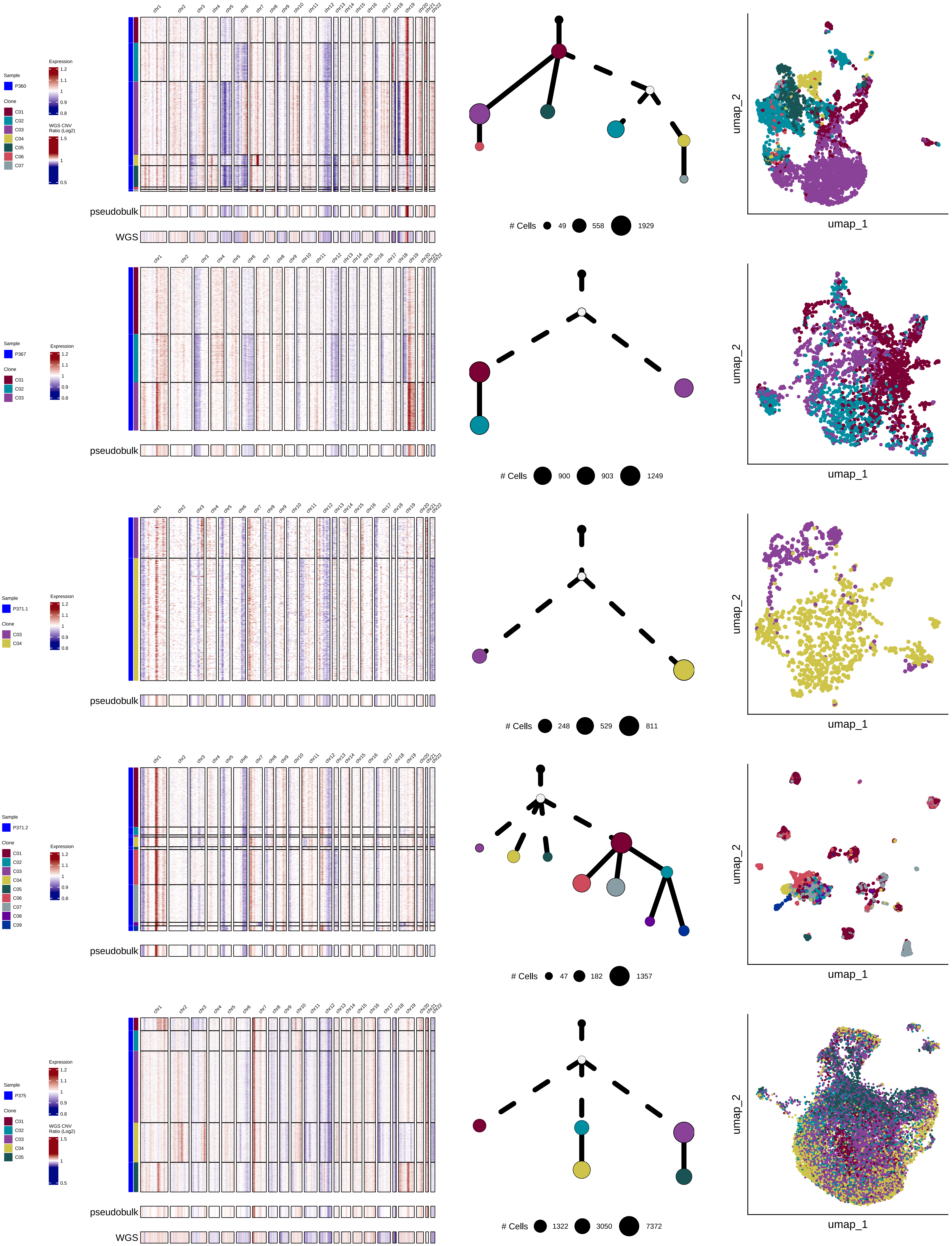

Sample
P360
Clone
C01
C02
C03
C04
C05
C06
C07
Expression
WGS CNV Ratio (Log2)
pseudobulk
WGS
### Cells
49
558
1929
umap_2
umap_1
Sample
P367
Clone
C01
C02
C03
Expression
pseudobulk
### Cells
900
903
1249
umap_2
umap_1
Sample
P371.1
Clone
C03
C04
Expression
pseudobulk
### Cells
248
529
811
umap_2
umap_1
Sample
P371.2
Clone
C01
C02
C03
C04
C05
C06
C07
C08
C09
Expression
pseudobulk
### Cells
47
182
1357
umap_2
umap_1
Sample
P375
Clone
C01
C02
C03
C04
C05
Expression
WGS CNV Ratio (Log2)
pseudobulk
WGS
### Cells
1322
3050
7372
umap_2
umap_1

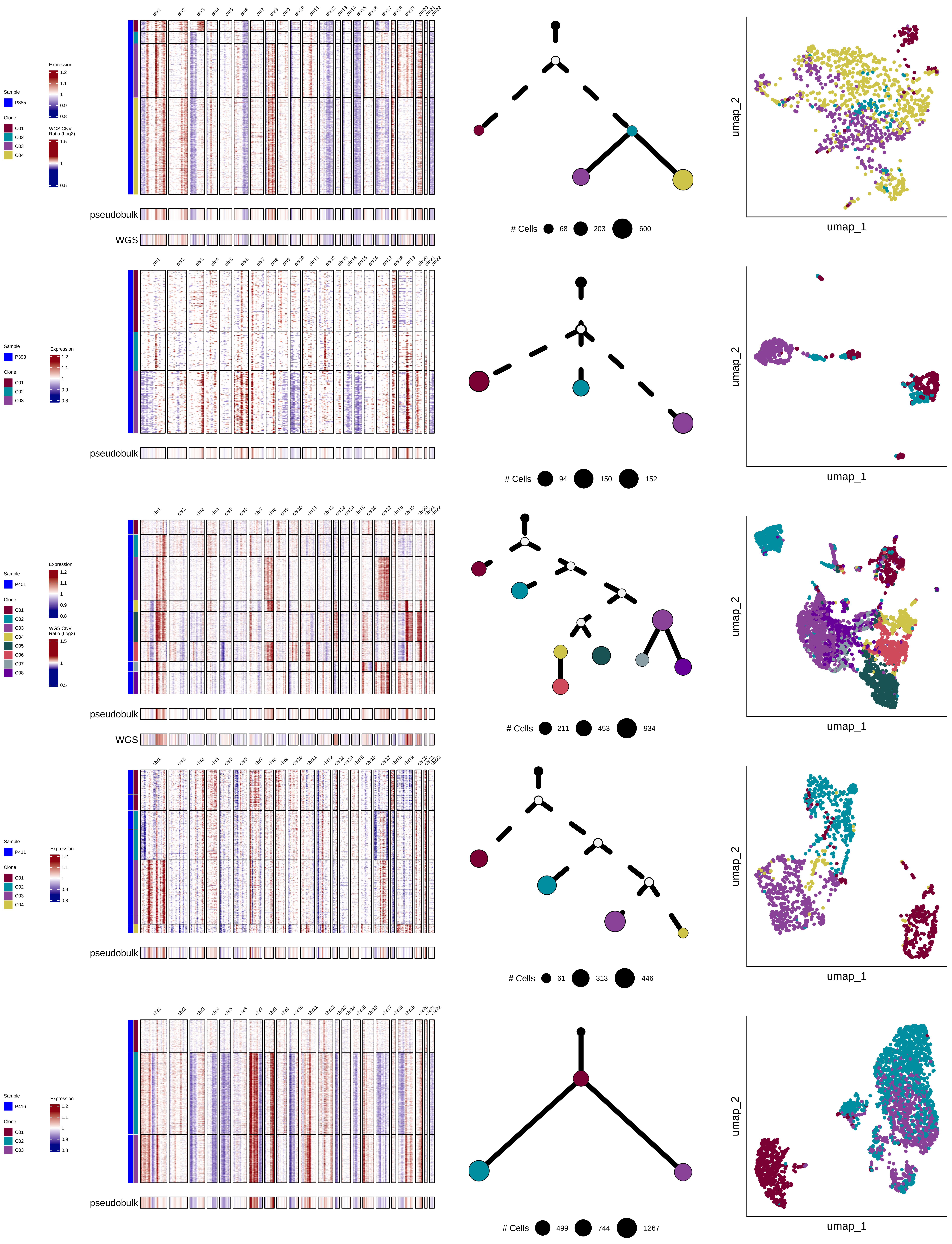

Sample
P385
Clone
C01
C02
C03
C04
Expression
1.2
1.1
1
0.9
0.8
WGS CNV Ratio (Log2)
1.5
1
0.5
chr1 chr2 chr3 chr4 chr5 chr6 chr7 chr8 chr9 chr10 chr11 chr12 chr13 chr14 chr15 chr16 chr17 chr18 chr19 chr20 chr21 chr22
pseudobulk
WGS
### Cells 68 203 600
umap_2
umap_1
Sample
P393
Clone
C01
C02
C03
Expression
1.2
1.1
1
0.9
0.8
pseudobulk
### Cells 94 150 152
umap_2
umap_1
Sample
P401
Clone
C01
C02
C03
C04
C05
C06
C07
C08
Expression
1.2
1.1
1
0.9
0.8
WGS CNV Ratio (Log2)
1.5
1
0.5
pseudobulk
WGS
### Cells 211 453 934
umap_2
umap_1
Sample
P411
Clone
C01
C02
C03
C04
Expression
1.2
1.1
1
0.9
0.8
pseudobulk
### Cells 61 313 446
umap_2
umap_1
Sample
P416
Clone
C01
C02
C03
Expression
1.2
1.1
1
0.9
0.8
pseudobulk
### Cells 499 744 1267
umap_2
umap_1

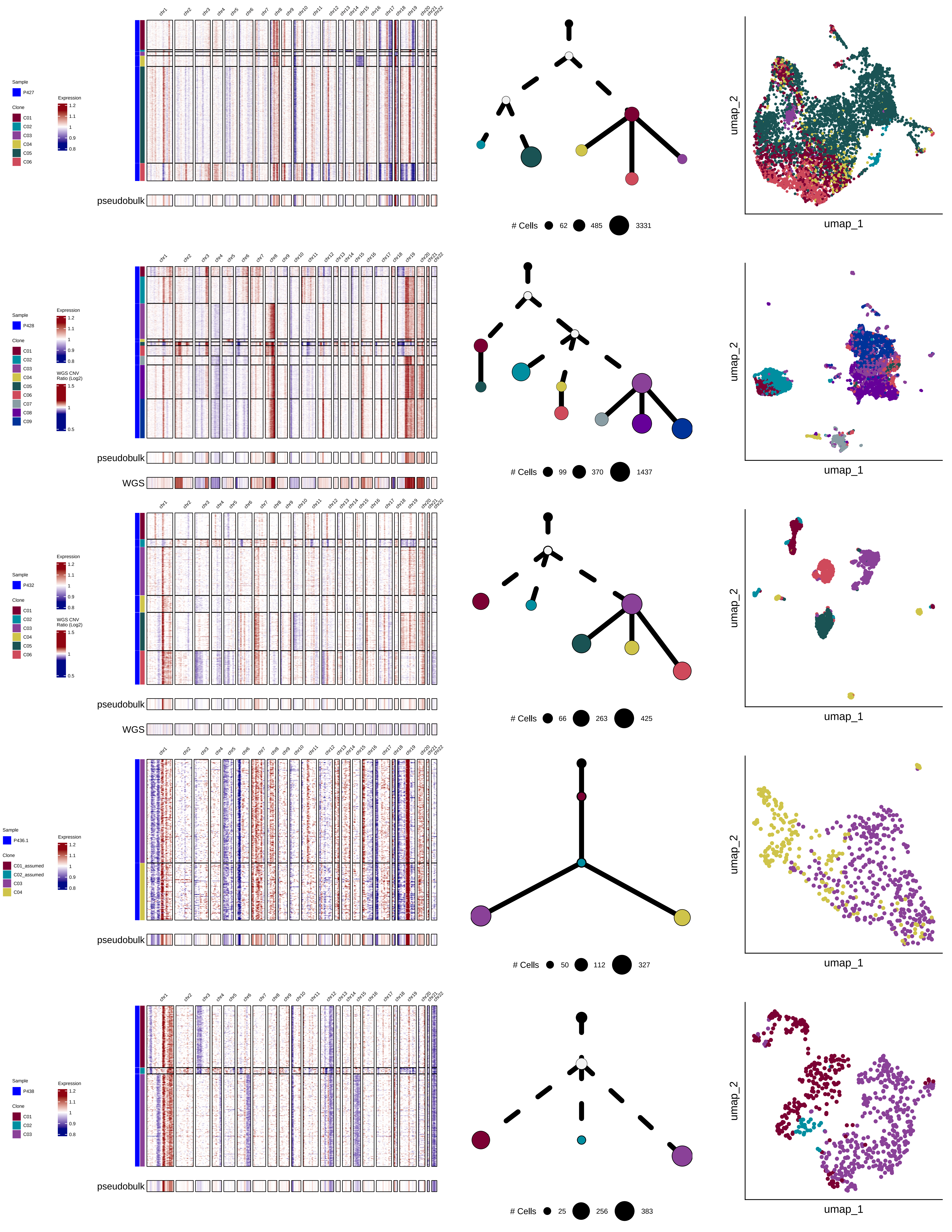

Sample
P427
Clone
C01
C02
C03
C04
C05
C06
Expression
1.2
1.1
1
0.9
0.8
pseudobulk
### Cells 62 485 3331
umap_2
umap_1
Sample
P428
Clone
C01
C02
C03
C04
C05
C06
C07
C08
C09
Expression
WGS CNV Ratio (Log2)
1.5
1
0.5
pseudobulk
WGS
### Cells 99 370 1437
Sample
P432
Clone
C01
C02
C03
C04
C05
C06
pseudobulk
WGS
### Cells 66 263 425
Sample
P436.1
Clone
C01_assumed
C02_assumed
C03
C04
pseudobulk
### Cells 50 112 327
Sample
P438
Clone
C01
C02
C03
pseudobulk
### Cells 25 256 383

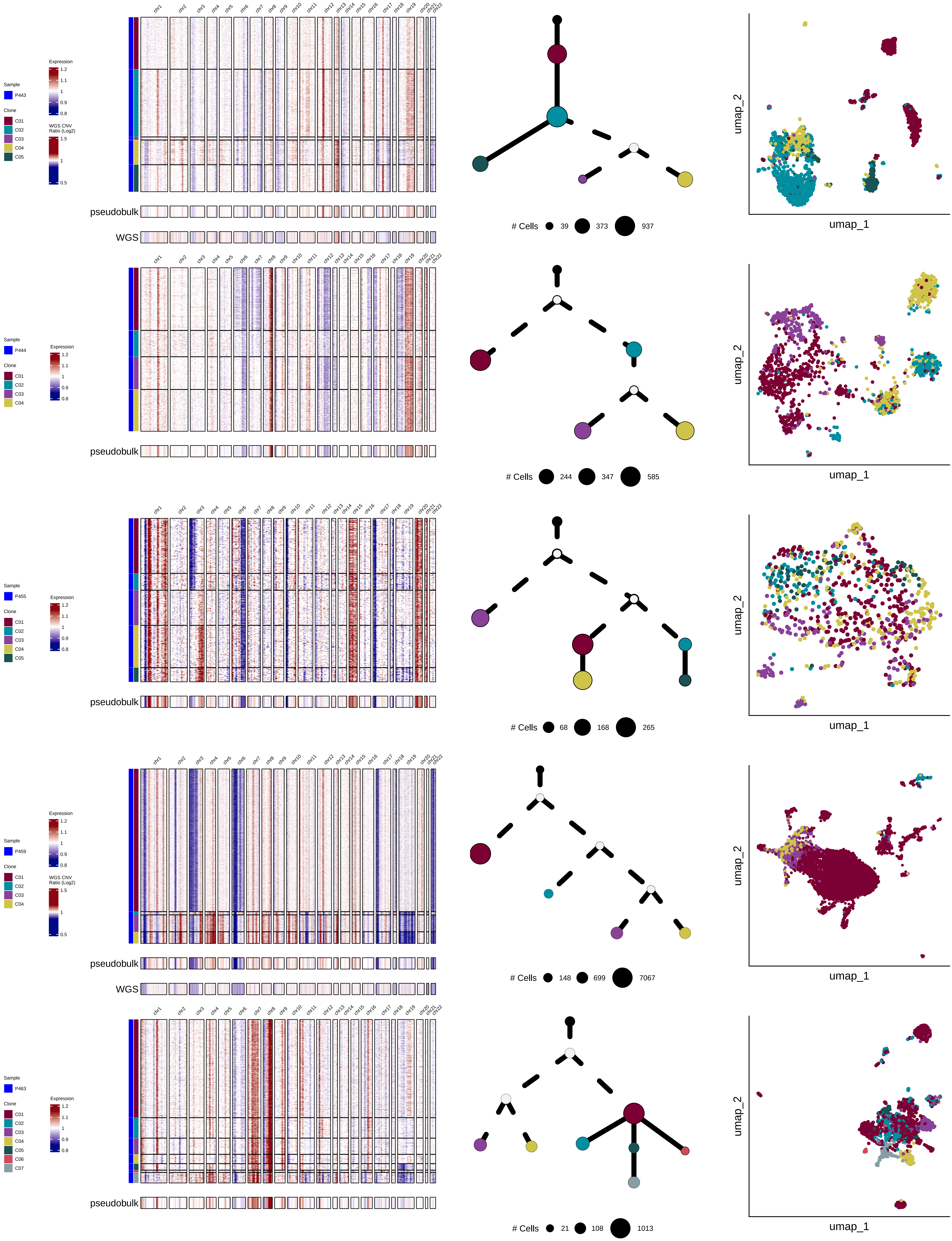

Sample
P443
Clone
C01
C02
C03
C04
C05
Expression
1.2
1.1
1
0.9
0.8
WGS CNV Ratio (Log2)
1.5
1
0.5
chr1 chr2 chr3 chr4 chr5 chr6 chr7 chr8 chr9 chr10 chr11 chr12 chr13 chr14 chr15 chr16 chr17 chr18 chr19 chr20 chr21 chr22
pseudobulk
WGS
### Cells 39 373 937
umap_2
umap_1
Sample
P444
Clone
C01
C02
C03
C04
Expression
1.2
1.1
1
0.9
0.8
pseudobulk
### Cells 244 347 585
umap_2
umap_1
Sample
P455
Clone
C01
C02
C03
C04
C05
Expression
1.2
1.1
1
0.9
0.8
pseudobulk
### Cells 68 168 265
umap_2
umap_1
Sample
P459
Clone
C01
C02
C03
C04
Expression
1.2
1.1
1
0.9
0.8
WGS CNV Ratio (Log2)
1.5
1
0.5
pseudobulk
WGS
### Cells 148 699 7067
umap_2
umap_1
Sample
P463
Clone
C01
C02
C03
C04
C05
C06
C07
Expression
1.2
1.1
1
0.9
0.8
pseudobulk
### Cells 21 108 1013
umap_2
umap_1

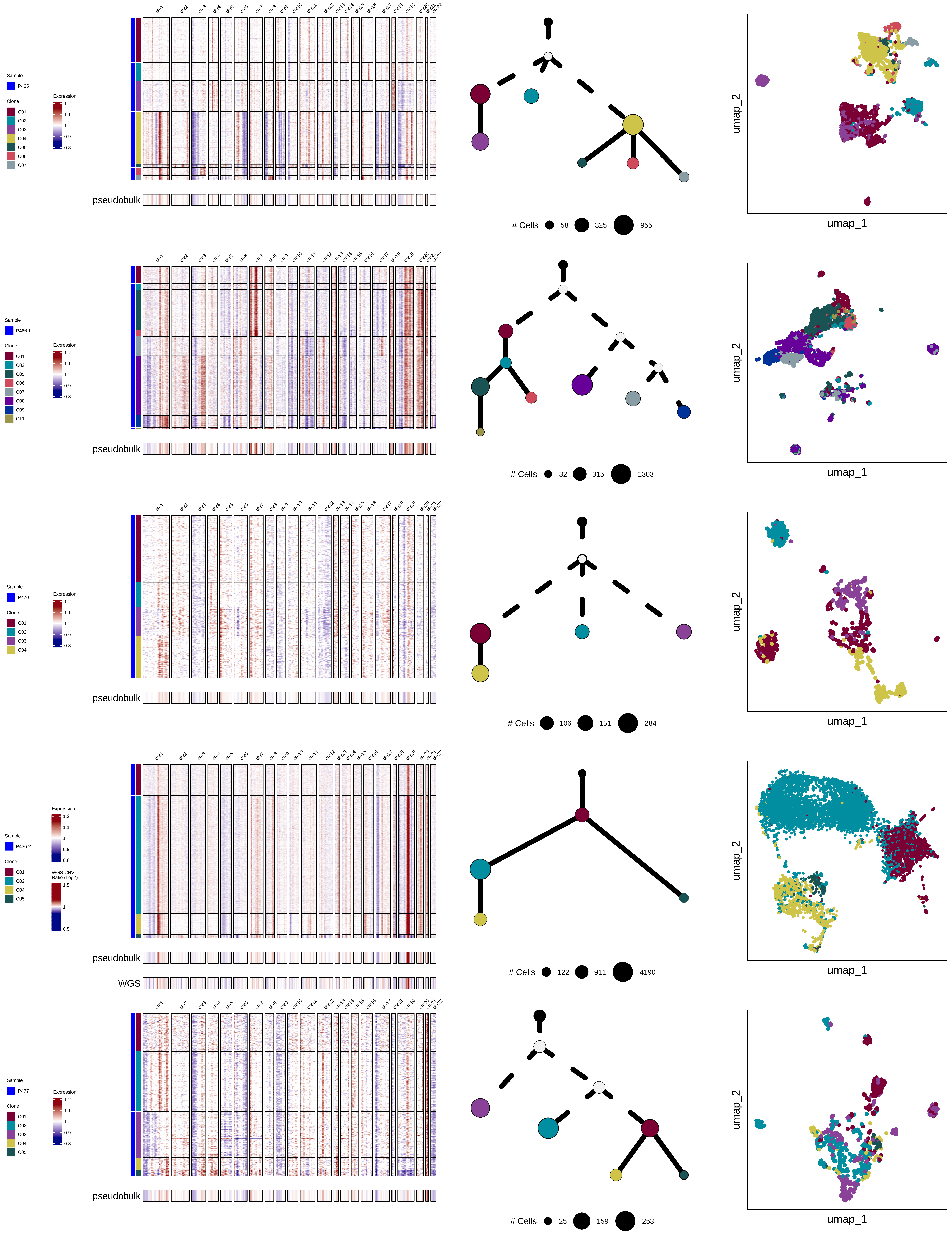
Sample
P465
Clone
C01
C02
C03
C04
C05
C06
C07
Expression
1.2
1.1
1
0.9
0.8
chr1
chr2
chr3
chr4
chr5
chr6
chr7
chr8
chr9
chr10
chr11
chr12
chr13
chr14
chr15
chr16
chr17
chr18
chr19
chr20
chr21
chr22
pseudobulk
### Cells
58
325
955
umap_2
umap_1
Sample
P466.1
Clone
C01
C02
C05
C06
C07
C08
C09
C11
Expression
1.2
1.1
1
0.9
0.8
pseudobulk
### Cells
32
315
1303
umap_2
umap_1
Sample
P470
Clone
C01
C02
C03
C04
Expression
1.2
1.1
1
0.9
0.8
pseudobulk
### Cells
106
151
284
umap_2
umap_1
Sample
P436.2
Clone
C01
C02
C04
C05
Expression
1.2
1.1
1
0.9
0.8
WGS CNV
Ratio (Log2)
1.5
1
0.5
pseudobulk
WGS
### Cells
122
911
4190
umap_2
umap_1
Sample
P477
Clone
C01
C02
C03
C04
C05
Expression
1.2
1.1
1
0.9
0.8
pseudobulk
### Cells
25
159
253
umap_2
umap_1

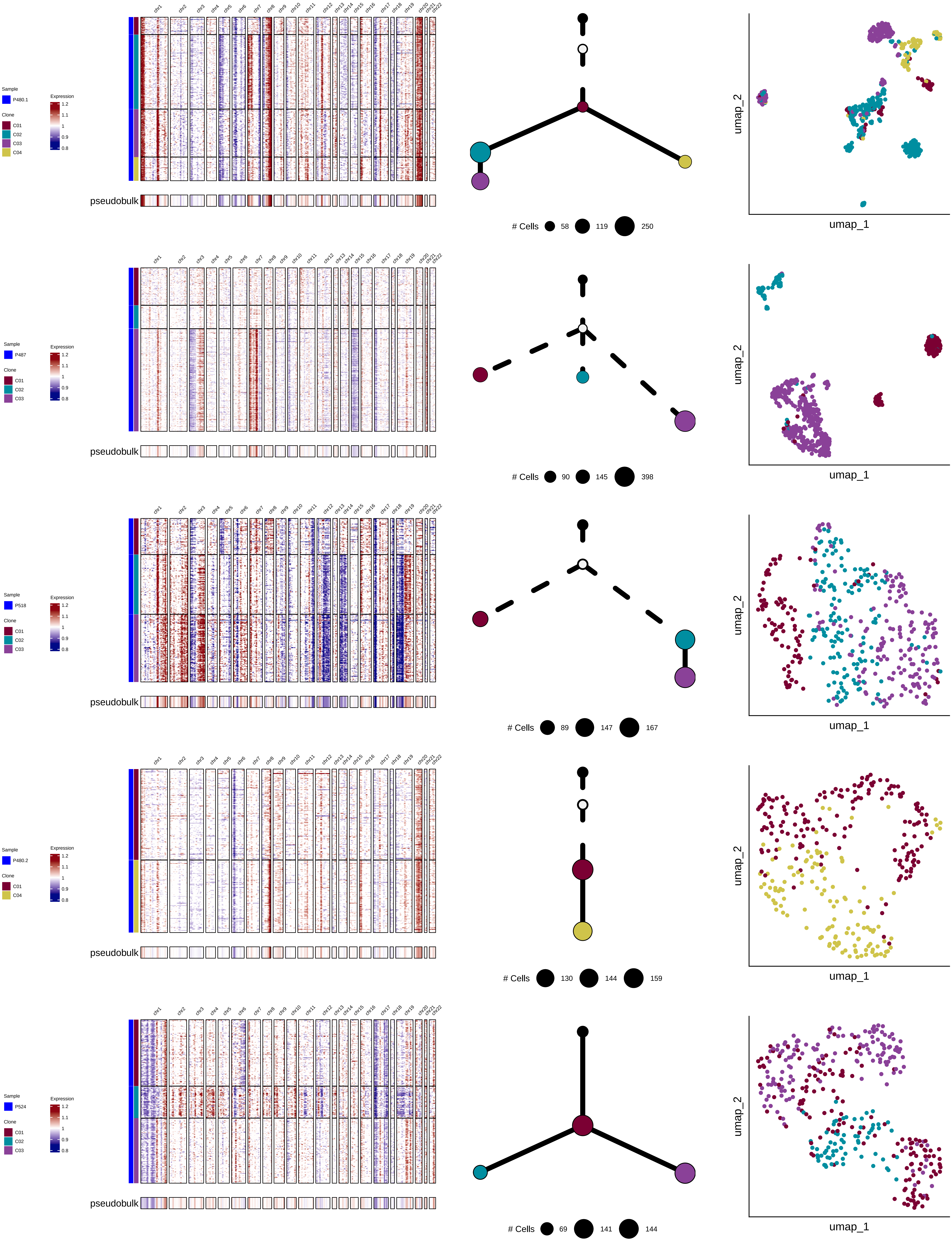
Sample
P480.1
Clone
C01
C02
C03
C04
Expression
1.2
1.1
1
0.9
0.8
chr1 chr2 chr3 chr4 chr5 chr6 chr7 chr8 chr9 chr10 chr11 chr12 chr13 chr14 chr15 chr16 chr17 chr18 chr19 chr20 chr21 chr22
pseudobulk
### Cells 58 119 250
umap_2
umap_1
Sample
P487
Clone
C01
C02
C03
Expression
1.2
1.1
1
0.9
0.8
pseudobulk
### Cells 90 145 398
umap_2
umap_1
Sample
P518
Clone
C01
C02
C03
Expression
1.2
1.1
1
0.9
0.8
pseudobulk
### Cells 89 147 167
umap_2
umap_1
Sample
P480.2
Clone
C01
C04
Expression
1.2
1.1
1
0.9
0.8
pseudobulk
### Cells 130 144 159
umap_2
umap_1
Sample
P524
Clone
C01
C02
C03
Expression
1.2
1.1
1
0.9
0.8
pseudobulk
### Cells 69 141 144
umap_2
umap_1

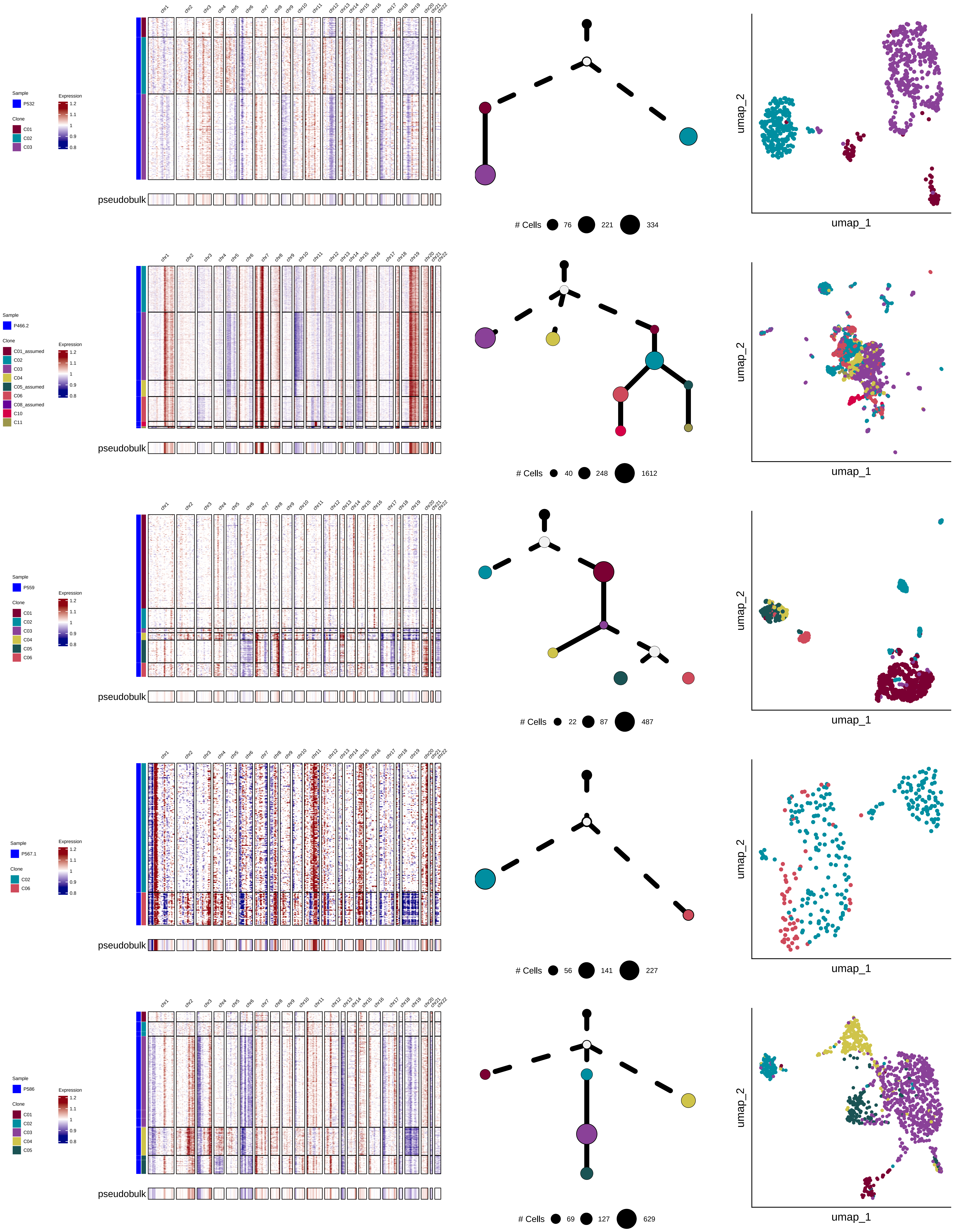

Sample
P532
Clone
C01
C02
C03
Expression
1.2
1.1
1
0.9
0.8
chr1 chr2 chr3 chr4 chr5 chr6 chr7 chr8 chr9 chr10 chr11 chr12 chr13 chr14 chr15 chr16 chr17 chr18 chr19 chr20 chr21 chr22
pseudobulk
### Cells 76 221 334
umap_2
umap_1
Sample
P466.2
Clone
C01_assumed
C02
C03
C04
C05_assumed
C06
C08_assumed
C10
C11
Expression
1.2
1.1
1
0.9
0.8
pseudobulk
### Cells 40 248 1612
umap_2
umap_1
Sample
P559
Clone
C01
C02
C03
C04
C05
C06
Expression
1.2
1.1
1
0.9
0.8
pseudobulk
### Cells 22 87 487
umap_2
umap_1
Sample
P567.1
Clone
C02
C06
Expression
1.2
1.1
1
0.9
0.8
pseudobulk
### Cells 56 141 227
umap_2
umap_1
Sample
P586
Clone
C01
C02
C03
C04
C05
Expression
1.2
1.1
1
0.9
0.8
pseudobulk
### Cells 69 127 629
umap_2
umap_1

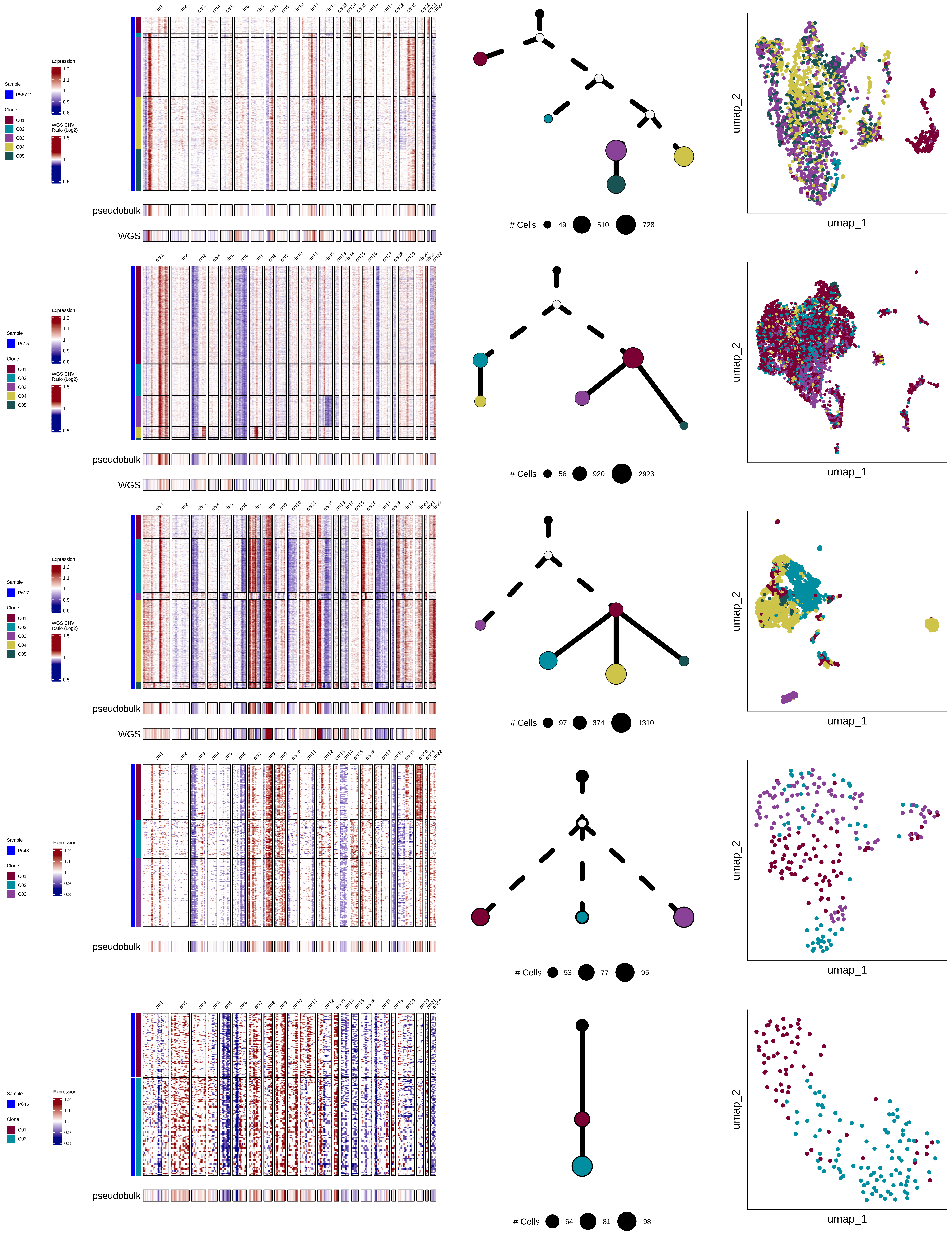

Sample
P567.2
P615
P617
P643
P645
Clone
C01
C02
C03
C04
C05
Expression
WGS CNV Ratio (Log2)
pseudobulk
WGS
### Cells
49
510
728
56
920
2923
97
374
1310
53
77
95
64
81
98
umap_1
umap_2

**Suppl. Figure 1. Clonal composition across all samples**
