## Supplementary material for "Clonal Heterogeneity in Human Pancreatic Ductal Adenocarcinoma and Its Impact on Tumor Progression": Suppl. Fig. 2

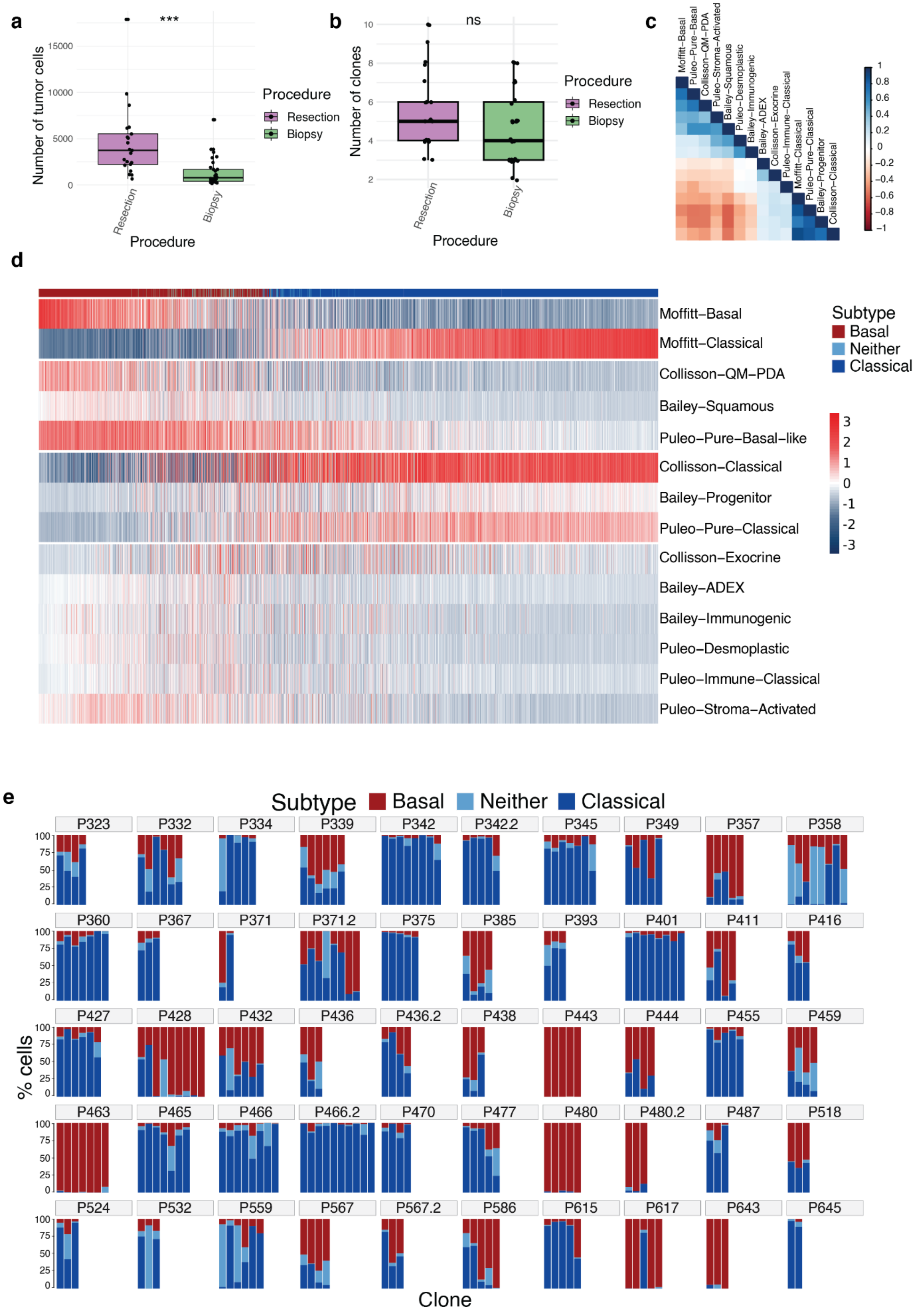
a
***
Number of tumor cells
15000
10000
5000
0
Resection
Biopsy
Procedure
Procedure
Resection
Biopsy
b
ns
Number of clones
10
8
6
4
2
Resection
Biopsy
Procedure
Procedure
Resection
Biopsy
c
Moffitt−Basal
Puleo−Pure−Basal
Collison−QM−PDA
Puleo−Stroma−Activated
Bailey−Squamous
Puleo−Desmoplastic
Bailey−Immunogenic
Bailey−ADEX
Collisson−Exocrine
Puleo−Immune−Classical
Moffitt−Classical
Puleo−Pure−Classical
Bailey−Progenitor
Collisson−Classical
1
0.8
0.6
0.4
0.2
0
−0.2
−0.4
−0.6
−0.8
−1
d
Moffitt−Basal
Moffitt−Classical
Collisson−QM−PDA
Bailey−Squamous
Puleo−Pure−Basal−like
Collisson−Classical
Bailey−Progenitor
Puleo−Pure−Classical
Collisson−Exocrine
Bailey−ADEX
Bailey−Immunogenic
Puleo−Desmoplastic
Puleo−Immune−Classical
Puleo−Stroma−Activated
Subtype
Basal
Neither
Classical
3
2
1
0
-1
-2
-3
e
Subtype
Basal
Neither
Classical
P323
P332
P334
P339
P342
P342.2
P345
P349
P357
P358
P360
P367
P371
P371.2
P375
P385
P393
P401
P411
P416
P427
P428
P432
P436
P436.2
P438
P443
P444
P455
P459
P463
P465
P466
P466.2
P470
P477
P480
P480.2
P487
P518
P524
P532
P559
P567
P567.2
P586
P615
P617
P643
P645
100
75
50
25
0
% cells
Clone

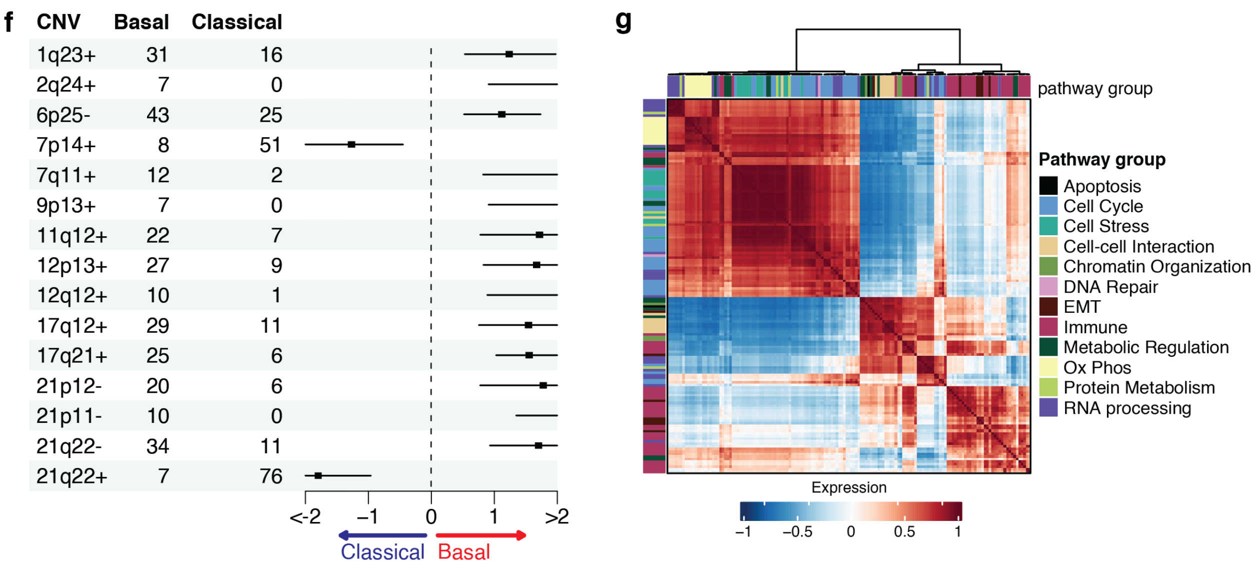


**Suppl. Figure 2.** a. Boxplot showing the number of tumor cells by procedure type. b. Boxplot showing the number of clones by procedure type. c. Correlation of published PDAC subtypes. d. Enrichment of published PDAC subtypes per cell. e. Percentage of cells expressing each subtype within each clone. f. Forest plot of significant CNVs between basal and classical clones. g. Correlation of pathway enrichment.
