## Supplementary material for "Clonal Heterogeneity in Human Pancreatic Ductal Adenocarcinoma and Its Impact on Tumor Progression": Suppl. Fig. 3

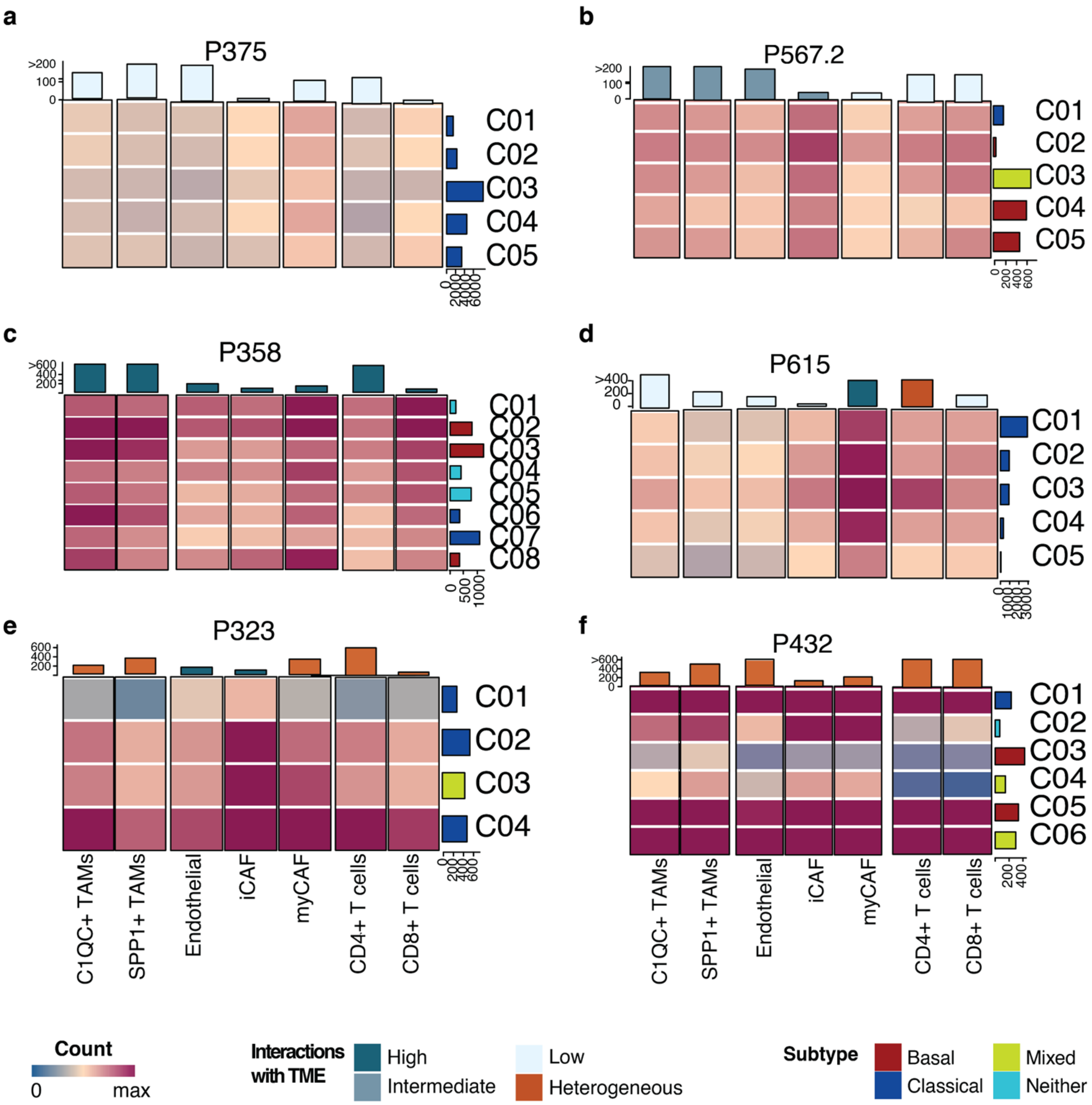


**Suppl. Figure 3**: a-f. Examples of samples with clones with comparable number of interactions with the TME (a-c), with clones interacting differently with some cell types (d-e), and with clones interacting differently with all TME cell types (f).
