## Supplementary material for "Clonal Heterogeneity in Human Pancreatic Ductal Adenocarcinoma and Its Impact on Tumor Progression": Suppl. Fig. 4

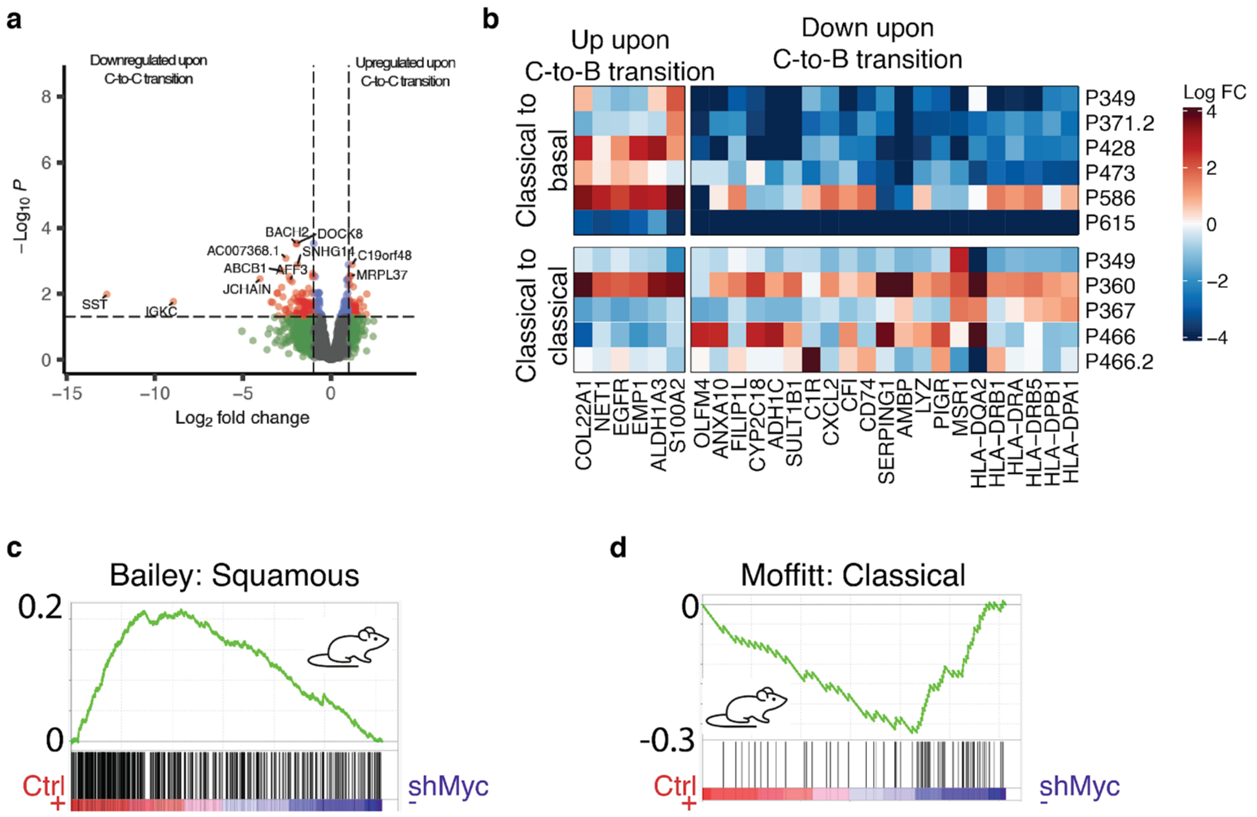


**Suppl. Figure 4:** a. Log fold change of selected gene expression between the pairs of classical to basal transition and classical to classical transition that include the 8q24 amplification. b. Genes differentially expressed upon the classical-to-classical evolutionary transition with 8q24 gain. c,d. *Myc* knockdown in two *Myc*+ basal mouse cell lines results in the depletion of the basal (c) and the enrichment of the classical subtype (d).
