## Supplementary material for "Clonal Heterogeneity in Human Pancreatic Ductal Adenocarcinoma and Its Impact on Tumor Progression": Suppl. Fig. 5

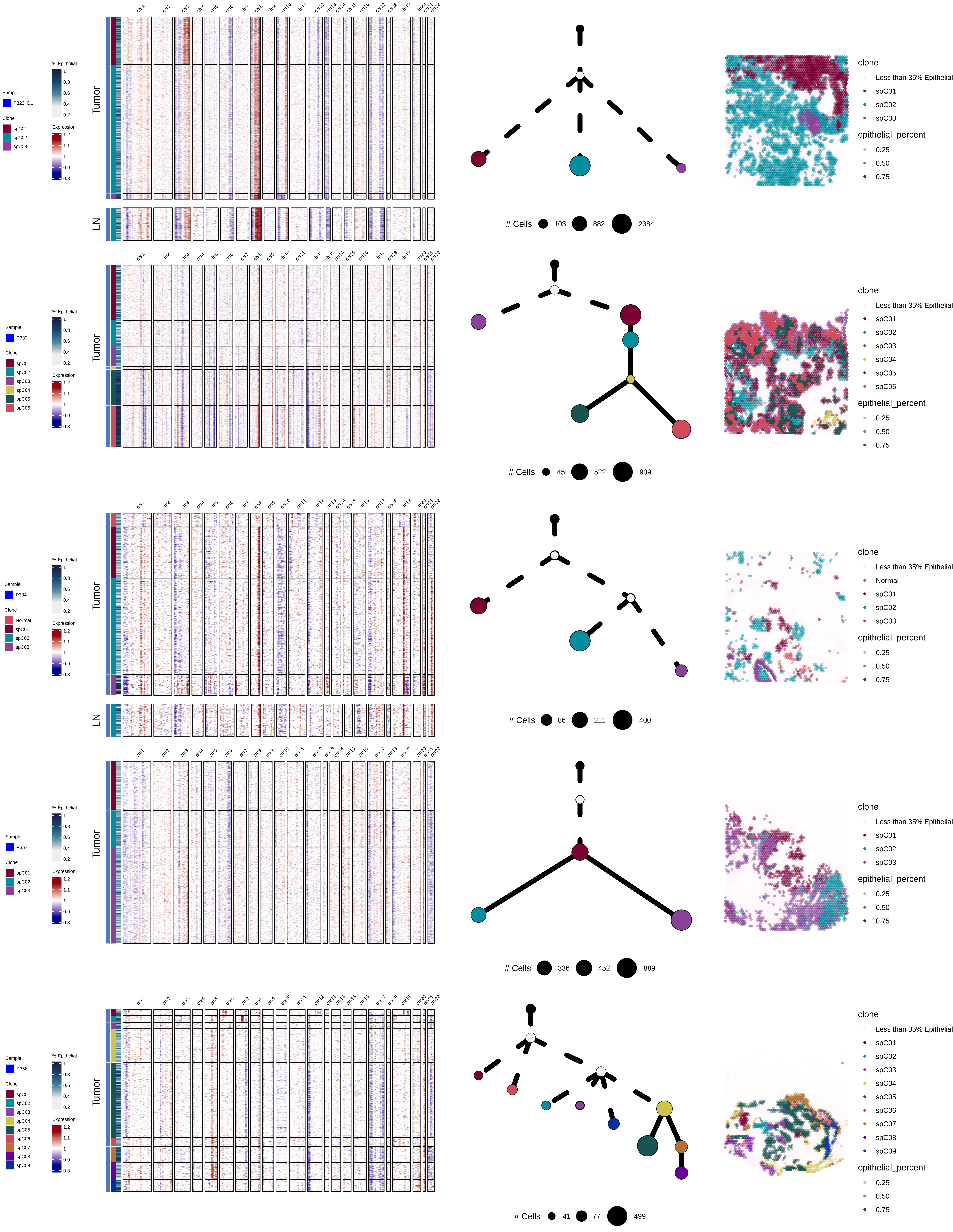
Sample
P323−D1
Clone
spC01
spC02
spC03
% Epithelial
Expression
Tumor
LN
chr1 chr2 chr3 chr4 chr5 chr6 chr7 chr8 chr9 chr10 chr11 chr12 chr13 chr14 chr15 chr16 chr17 chr18 chr19 chr20 chr21 chr22
### Cells 103 882 2384
clone
Less than 35% Epithelial
epithelial_percent
0.25
0.50
0.75
P332
spC04
spC05
spC06
### Cells 45 522 939
P334
Normal
### Cells 86 211 400
P357
### Cells 336 452 889
P358
spC07
spC08
spC09
### Cells 41 77 499

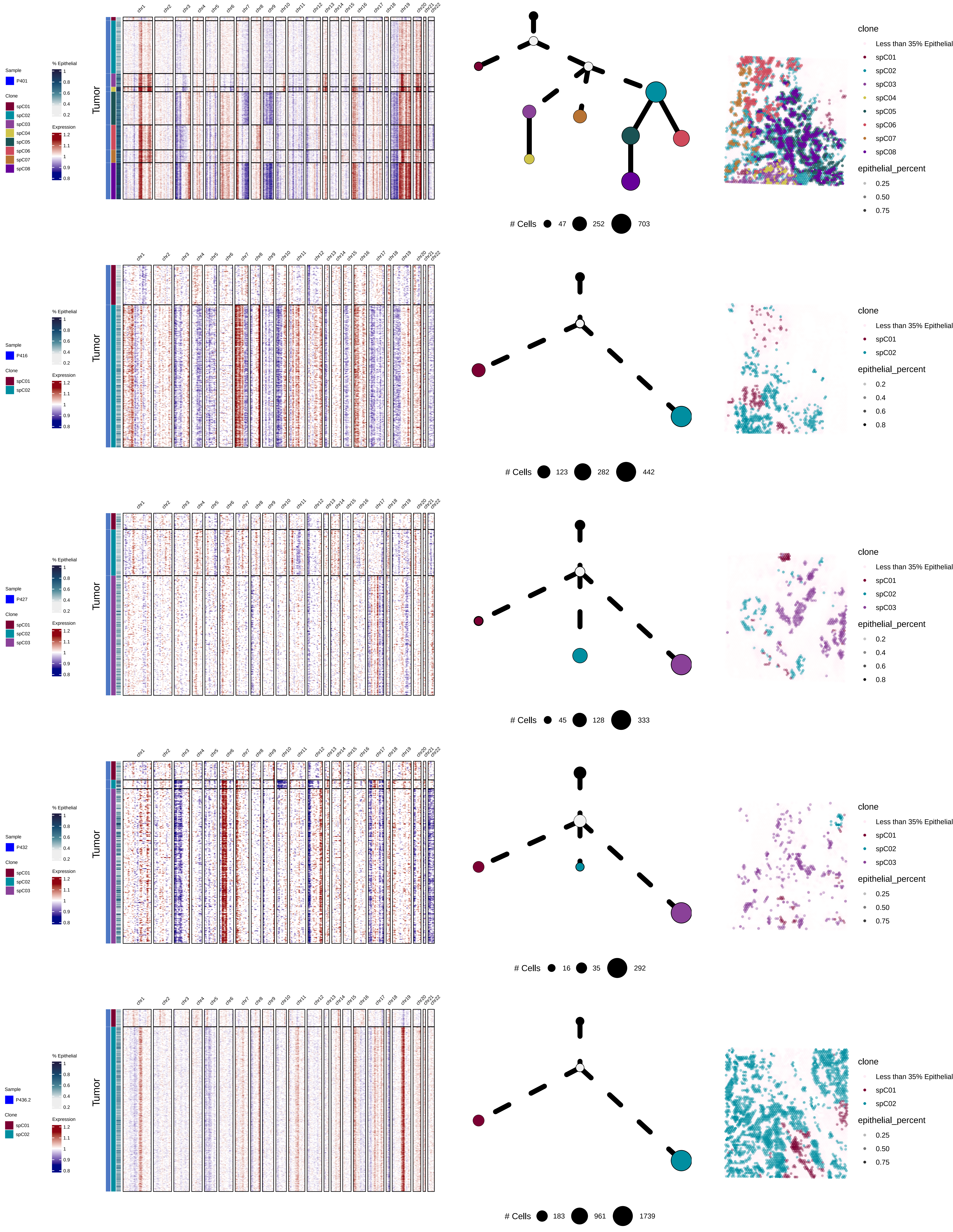

Sample
P401
Clone
spC01
spC02
spC03
spC04
spC05
spC06
spC07
spC08
% Epithelial
1
0.8
0.6
0.4
0.2
Expression
1.2
1.1
1
0.9
0.8
Tumor
chr1 chr2 chr3 chr4 chr5 chr6 chr7 chr8 chr9 chr10 chr11 chr12 chr13 chr14 chr15 chr16 chr17 chr18 chr19 chr20 chr21 chr22
### Cells 47 252 703
clone
Less than 35% Epithelial
spC01
spC02
spC03
spC04
spC05
spC06
spC07
spC08
epithelial_percent
0.25
0.50
0.75
Sample
P416
Clone
spC01
spC02
% Epithelial
1
0.8
0.6
0.4
0.2
Expression
1.2
1.1
1
0.9
0.8
Tumor
chr1 chr2 chr3 chr4 chr5 chr6 chr7 chr8 chr9 chr10 chr11 chr12 chr13 chr14 chr15 chr16 chr17 chr18 chr19 chr20 chr21 chr22
### Cells 123 282 442
clone
Less than 35% Epithelial
spC01
spC02
epithelial_percent
0.2
0.4
0.6
0.8
Sample
P427
Clone
spC01
spC02
spC03
% Epithelial
1
0.8
0.6
0.4
0.2
Expression
1.2
1.1
1
0.9
0.8
Tumor
chr1 chr2 chr3 chr4 chr5 chr6 chr7 chr8 chr9 chr10 chr11 chr12 chr13 chr14 chr15 chr16 chr17 chr18 chr19 chr20 chr21 chr22
### Cells 45 128 333
clone
Less than 35% Epithelial
spC01
spC02
spC03
epithelial_percent
0.2
0.4
0.6
0.8
Sample
P432
Clone
spC01
spC02
spC03
% Epithelial
1
0.8
0.6
0.4
0.2
Expression
1.2
1.1
1
0.9
0.8
Tumor
chr1 chr2 chr3 chr4 chr5 chr6 chr7 chr8 chr9 chr10 chr11 chr12 chr13 chr14 chr15 chr16 chr17 chr18 chr19 chr20 chr21 chr22
### Cells 16 35 292
clone
Less than 35% Epithelial
spC01
spC02
spC03
epithelial_percent
0.25
0.50
0.75
Sample
P436.2
Clone
spC01
spC02
% Epithelial
1
0.8
0.6
0.4
0.2
Expression
1.2
1.1
1
0.9
0.8
Tumor
chr1 chr2 chr3 chr4 chr5 chr6 chr7 chr8 chr9 chr10 chr11 chr12 chr13 chr14 chr15 chr16 chr17 chr18 chr19 chr20 chr21 chr22
### Cells 183 961 1739
clone
Less than 35% Epithelial
spC01
spC02
epithelial_percent
0.25
0.50
0.75

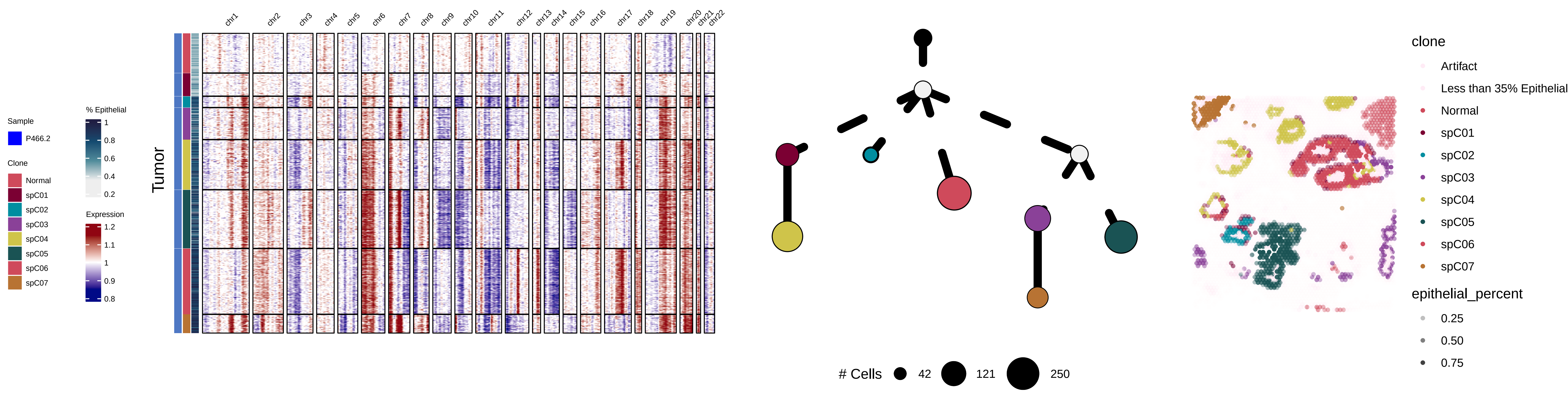


**Suppl. Figure 5. Clonal composition  and distribution across space for all samples profiled with spatial transcriptomics.**
