## Supplementary material for "Clonal Heterogeneity in Human Pancreatic Ductal Adenocarcinoma and Its Impact on Tumor Progression": Suppl. Fig. 6

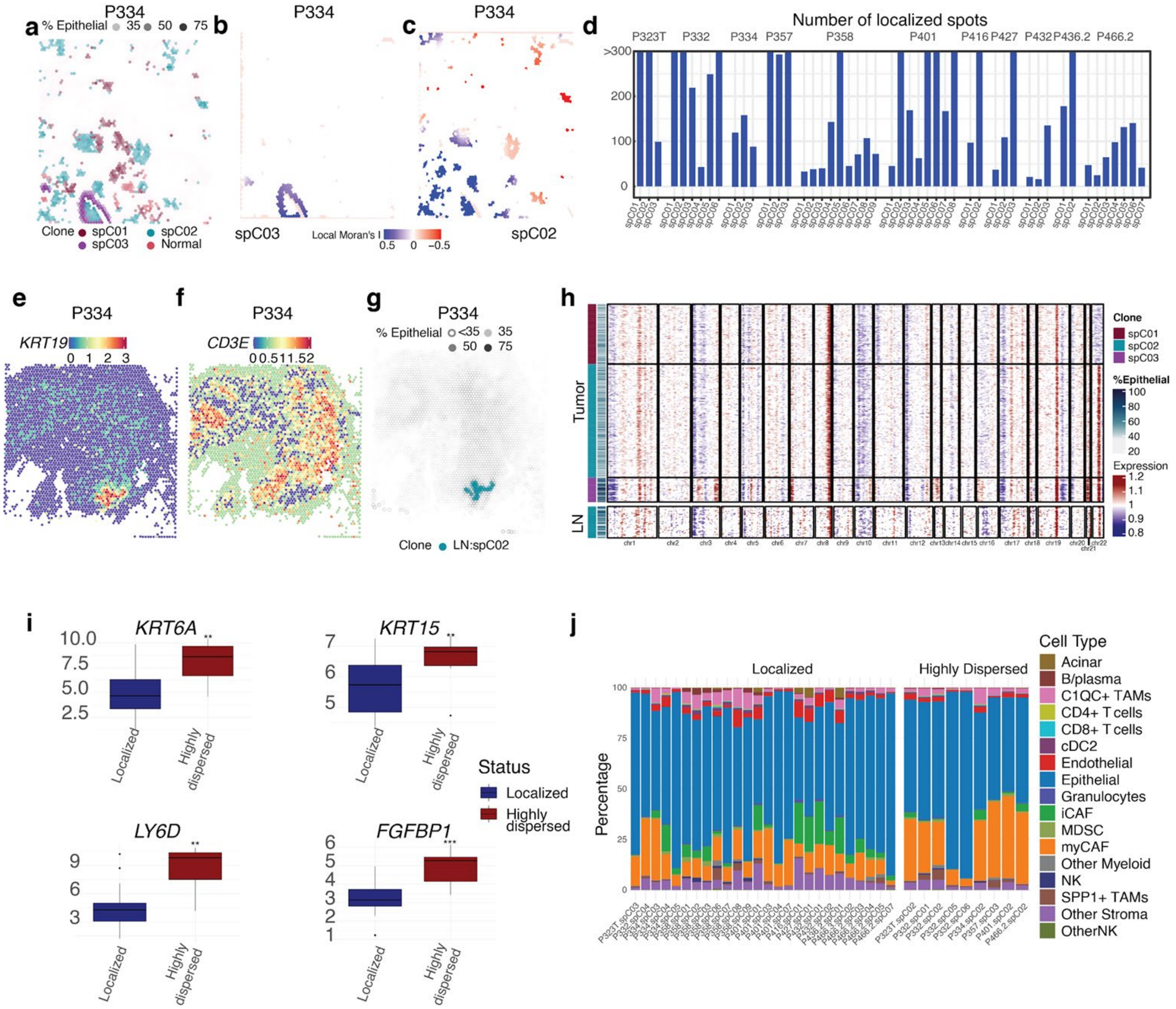


**Suppl. Fig. 6. Dispersed clonal populations facilitate lymph node invasion in PDAC**: a. Spatial mapping of the clones in the profiled region of a representative sample (P334). b. Local Moran's I score for clone spCNV2 of P334. c. Local Moran's I score for clone spCNV3 of P334. d. Barplot showing the number of localized spots for each clone of each sample. e. *KRT19* expression in the lymph node of P334. f. *CD3E* expression in the lymph node of P334. g. Spatial mapping of spCNV2 clone that has invaded P334's lymph node. h. CNV profile of spots of the tumor region (top) and the lymph node (bottom) of 334 with more than 35% epithelial content. Clustered spots are considered clones. i. Boxplots of basal marker gene expression in highly dispersed and localized clones. j. Cell type composition in localized and highly dispersed clones.
